# Protracted development of visuo-proprioceptive integration for uni-and bimanual motor coordination

**DOI:** 10.1101/601435

**Authors:** Marie Martel, Jose P. Ossandón, Boukje Habets, Tobias Heed

## Abstract

Sensory and motor processes undergo massive developmental changes over at least a decade in human development. Interdependencies between different sensorimotor control mechanisms as well as bodily abilities are difficult to assess when isolated experiments are tested in small age ranges. Here, we assessed coordinative abilities of 120 children aged 4-12 for the two hands or a hand with another sensory signal in multiple, highly comparable sensorimotor tasks. This multi-task approach allowed assessing the development and interplay of several aspects of motor control related to different coordinative requirements. Children were first able to symmetrically move the two hands, and only later to coordinate one hand with a proprioceptive or visual signal. The ability to strategically ignore sensory information was available last. The pattern of partial correlations among tasks suggests protracted, interdependent, chained development within individuals.

**NEW AND NOTEWORTHY:** Development unfolds as a cascade: each new ability sets the stage for learning further skills in motor, sensory, cognitive, and social domains. Here, we charted the performance of 4–12- year-olds in six coordinative tasks that are all based on a common experimental paradigm but address three different sensorimotor-cognitive domains. This approach characterizes dependencies between multiple aspects of cognitive modulation in the interplay of sensory integration and motor control.

## 1. INTRODUCTION

Human motor development unfolds over two decades. New motor skills often change how children perceive and interact with their environment. This interplay sets the stage for learning further skills in sensorimotor, cognitive, and social domains (Adolph & Franchak, 2017; Adolph & Hoch, 2019; Hamilton et al., 2016). Unsurprisingly, then, sensory development, too, extends over long timescales. For instance, children’s multisensory processing differs markedly from that of adults until at least 8-10 years of age (for review, see Burr & Gori, 2012; Gori, 2015). According to current theories of motor control, perception and movement are linked by internal models – processes that predict the sensory consequences of planned and ongoing movement to enable motor corrections when expected and true sensory consequences differ (Haar & Donchin, 2020; Shadmehr & Krakauer, 2008). However, knowledge about how the respective sensorimotor interplay and cognitive organization develop is fractured.

Developmental research requires extensive effort. Many studies have, therefore, focused on limited age ranges as well as on only one or few sensorimotor functions or tasks. This research strategy makes it difficult to characterize developmental trajectories and to determine how specific abilities build on others. Therefore, the protracted and interwoven nature of sensorimotor development calls for studies that investigate wide age ranges and multiple sensorimotor functions. As one step in this direction, we present here a set of 6 sensorimotor coordination tasks that were performed by 120 children aged 4-12 years as well an adult group. We designed all tasks based on a common experimental framework in which one or both hands had to be moved in circles. The study’s rationale is illustrated in Fig. 1A, B. We tested children’s ability to coordinate symmetrically with one and two hands, as well as asymmetrically. An overarching task design allows us to characterize, with three task pairs, the development of, and dependencies between, four distinct aspects of sensorimotor coordination (see Fig. 1B). We briefly introduce these different aspects here.

**Fig. 1:**
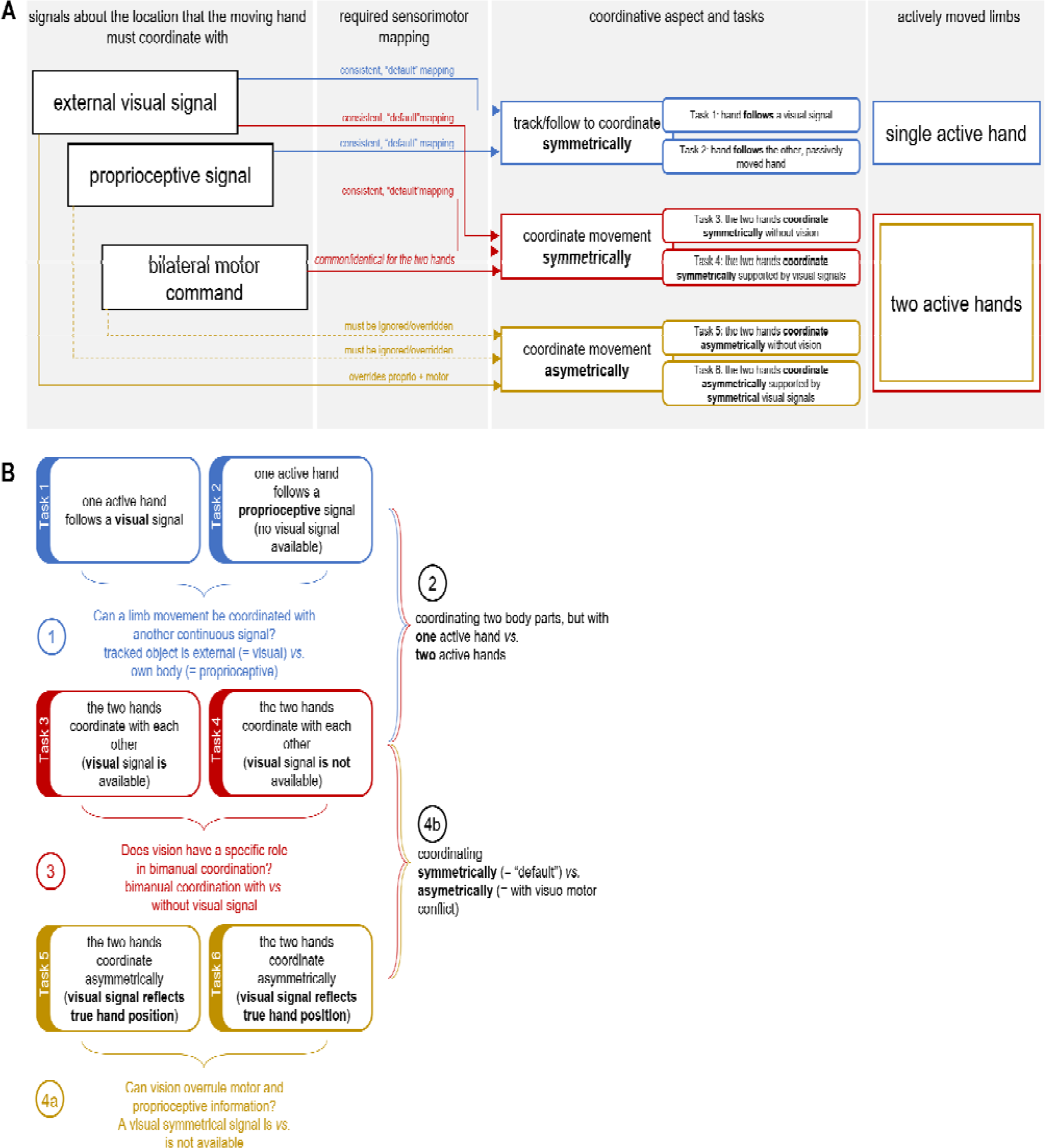
Study rationale. The illustration in (**A)** contrasts the six tasks according to relevant sensory signals, required sensorimotor mapping, coordination, and involved hands. Six tasks, grouped into three pairs, tested three distinct aspects of sensorimotor hand coordination. Two tasks required actively moving a single hand, whereas the remaining four tasks were bimanual (right-most panel, “actively moved limbs”). The different tasks provided different signals about the active hand’s location (left-most panel, “signals about the location…”): depending on the task, hand location could be sensed only proprioceptively or proprioceptively and visually. In bimanual tasks, coordination could further be affected by motor signals being identical or different for the two hands. Depending on the the task (third panel, “coordinative aspect and tasks”), locations either mapped consistently between the different sensory/motor modalities (solid arrows; arrow colors match those of the task pairs for overview) or, instead, the location signaled by some modality had to be ignored to let a different sensory signal dominate coordination (dashed arrows). The color of the mapping arrows matches with the tasks. Note, that the first, unimanual and the second, bimanual task pairs are based on only consistent location information. They differ in that the first pair does not afford a bilateral motor signal to base coordination on. The third task pair differs from the second in requiring suppression of inconsistent information. The illustration in **(B)** contrasts the four aspects, indicated by the numbers in circles, of motor coordination afforded by the different comparisons between tasks and task sets, respectively. The two tasks of each task set, as well as comparison between the different tasks sets, each address specific research questions. Numbering 4a and 4b indicates that these two comparisons speak to the same underlying coordinative aspect.

The first aspect (Fig. 1A, B: Tasks 1 & 2, blue shapes, Fig. 1B: circle number 1) is coordinating a hand movement with a sensory signal. Participants had to guide their dominant hand’s position continuously via a visual or passive-proprioceptive signal. For visual tracking (Task 1), a cursor moved in circles on one side of the display, and participants had to match a moving cursor that represented their dominant hand as closely as possible. For passive-proprioceptive tracking (Task 2), we moved the participant’s non-dominant hand in a circle, and they had to match its position continually with their dominant hand. Thus, the two tasks compare coordination with an external, non-body object vs. the own body.

This type of tracking task is a basic implementation of a sensorimotor loop. Previous research has addressed such tasks in adults and children. Adults successfully track both visual and proprioceptive information (Buekers et al., 2000; Jones et al., 2010; Tanaka et al., 2009). Children aged 5-7 were able to perform visual tracking, as investigated in the present study (Gauthier et al., 1988). However, improvement was evident until age 10-11 (Van Roon et al., 2008) or even late adolescence (Zanone, 1990). Many developmental studies have observed step-like performance improvements and temporary performance decrements within circumscribed time intervals (e.g. Badan et al., 2000; Hay, 1978; Martel et al., 2020; Pellizzer & Hauert, 1996; Smyth et al., 2004), suggesting that development is non-linear. We included the two simple tracking tasks as the baseline against which we can compare the remaining tasks. Moreover, tracking has not been characterized across the entire childhood range (∼4-12 years, i.e., up to about puberty).

The second aspect we consider (Fig. 1A, B: Tasks 3 & 4, red shapes, Fig. 1B: circle number 2) is the coordination of two actively moving body parts. Participants executed symmetrical movements with the two hands. Compared to the first task pair (i.e., blue vs. red shapes in Fig. 1), in which only a single hand was actively moved, bimanual movement affords a third signal that may help coordination: the two hands can both be driven by a common motor command, which would “automatically” create symmetry. As a third aspect (Fig. 1B: circle number 3), we tested coordination both with (Task 3) and without vision (Task 4). Without vision, the task directly compares to unimanual tracking of the own, passively moved hand, with the difference that now both hands move actively. Adding a visual signal allows testing whether this additional source can aid coordination over and above motor and proprioceptive information.

We presume that easier functionalities develop earlier than more difficult ones. There are two possible outcomes regarding the additional requirement of actively moving the second hand: an additional signal might increase coordinative difficulty. Alternatively, the common motor signal for the two limbs may provide an easy-to-use mechanism for coordination, making bimanual easier than unimanual coordination.

Previous research clearly favors the second alternative. Adults preferentially synchronize bimanual movements symmetrically, especially when movement speed is high (e.g. Brandes, Rezvani, & Heed, 2017; Kelso, Scholz, & Schöner, 1986; Spencer, Ivry, Cattaert, & Semjen, 2005; for review, see Gooijers & Swinnen, 2014). Young children usually (de Boer et al., 2012; Fagard et al., 1985; Jeeves et al., 1988; Lantero & Ringenbach, 2007), though not always (Fagard, 1987), exhibit lower accuracy for bimanual symmetry than older children and adults. Nonetheless, they often coordinate the two hands symmetrically involuntarily (Cincotta & Ziemann, 2008; Lazarus & Todor, 1987), a phenomenon that declines around 10 years of age and has been associated with the maturation of the corpus callosum (Cincotta & Ziemann, 2008; de Boer et al., 2012) but may also involve cerebellar development (Boisgontier et al., 2018). As a further developmental aspect, the use of visual information to aid (bimanual) motor coordination may develop rather late. For instance, children aged 4 and 6 disregarded visual information during circle drawing, contrary to 8 year-olds and adults (Lantero & Ringenbach, 2007). Such results suggest critical developmental changes in sensorimotor integration during this age range. Therefore, we included two bimanual coordination tasks that are directly comparable to our unimanual tracking tasks – one that does, and one that does not include visual information of the two hands’ current position.

The fourth aspect we consider (Fig. 1A, B: Tasks 5 & 6, ochre shapes) addresses complex sensorimotor coordination in which some sensory information is incongruent, so that some information must be selected but other information ignored to coordinate successfully. This requirement contrasts with a default situation, in which all sensory information is congruent and can potentially aid performance (Fig. 1B, circles number 4a, 4b). Even if the preference for symmetry in bimanual coordination may in part reflect synchronized motor commands (Cohen, 1971; Kelso, 1984; Riek & Woolley, 2005), many studies have demonstrated a pivotal role of symmetry or, more generally speaking, congruency of perceptual signals for effective motor coordination (Bingham, 2004; Brandes et al., 2017; Heed & Röder, 2014; Mechsner et al., 2001; for review, see Shea et al., 2016). Regarding our circling tasks, it is impossible for most adults to circle the two hands at different speeds, such as doing four turns with the left for every three turns with the right hand). Critically, this task becomes manageable when the different hand speeds are transformed into same-speed visual signals, akin to turning handles connected to cogwheels of different size (Mechsner et al., 2001); the required, asymmetrical movement pattern is achieved by symmetrically synchronizing the visual cues.

To our knowledge, this asymmetric circling task has not been studied in children. However, developmental studies have employed designs similar to the “Etch-A-Sketch” drawing toy (Fagard, 1987; Fagard et al., 1985; Jeeves et al., 1988; Preilowski, 1972, 1975; Swinnen, 2002; see https://en.wikipedia.org/wiki/Etch_A_Sketch), also termed “Lissajous” display (Chiou & Chang, 2016; Kovacs et al., 2009). Each hand can manipulate a knob or handle; one hand’s knob controls a cursor’s horizontal, and the other its vertical position. When each knob is turned at constant speed, the cursor will draw a straight line, the slant of which depends on the two knobs’ speeds. Adults can maintain different knob-turning speeds with the two hands with, but not without the visual aid of the cursor. Children performed such tasks successfully at around 8-10 years of age –significantly later than the tasks we introduced earlier (Fagard, 1987; Fagard et al., 1985; Jeeves et al., 1988; Leinen et al., 2016).

Here, we included asymmetric, bimanual hand circling with and without visual, transformed cursor positions as tasks that are, again, directly comparable to the four other tasks we devised. We expect that participants will be unable to perform asymmetrically when the cursors reflect (asymmetric) true, untransformed hand positions and this task is included merely as a control task. The transformed cursor task has in common with unimanual coordination that it requires coordination with an external signal that does not directly reflect limb position. However, in contrast to unimanual coordination, participants must downweigh their proprioceptive input and overcome the motor system’s preference for symmetry. They must prioritize visual, symmetric information to coordinate asymmetric circling.

In sum, the present study related the development of three aspects of motor control: unimanual coordination with an external visual or passive-proprioceptive guiding signal; visual and proprioceptive coordination of active, symmetrical, bimanual control; and the ability to overcome the motor system’s symmetry preference by focusing on visual, and disregarding incongruent proprioceptive, information. Employing a within-subject design allows assessing whether functions that emerge later rely on development of those functions that emerge earlier by correlating performance across tasks.

If movement coordination were perceptually driven (Bingham, 2004), then bimanual coordination would be a subcategory of a more general ability to coordinate motorically by means of perception. Uni-and bimanual coordination should then show similar development. In contrast, if motor synchronization played a critical role in bimanual coordination, then one might expect different development for bimanual and unimanual tasks, with bimanual emerging earlier than unimanual coordination. Given that previous research suggests such visual-motor integration to emerge rather late, visually guided, asymmetrical coordination should also emerge late.

## 2. METHODS

### 2.1. Participants and ethical approval

We enrolled children at public science popularization events, school visits to our lab, and via flyers, until we had recruited at least 12 children of each age from 4-12 years that met our inclusion criteria. We did not include older ages because pubertal development is known to affect motor performance (e.g. Martel et al., 2021; Viel et al., 2009), so that we would have had to assess puberty on top of age, which seemed inappropriate for most of the younger children. We included 120 (52 girls, 68 boys) of 138 tested children in the final sample. They had normal or corrected-to-normal vision and were free of any known neurological, visual, and tactile impairments and disorders. They were born in regular term (> 37^th^ gestational week), scored between 70 and 130 on a short IQ test (3 children excluded), and passed screening for attentional disorders, learning disabilities and delayed psychomotor acquisition (11 children excluded; see supplementary information at https://osf.io/grnbx for details). We determined hand dominance with the quick oral version of the Edinburgh scale, asking participants their hand preference for writing, drawing, and using a scissor or spoon (Oldfield, 1971). We let younger children handle a pair of scissors and briefly write/draw something to assess their hand dominance directly and additionally confirmed our observations with the parents. The last 4 excluded children, all 4-year-old, were unable to follow the task instructions or did not want to complete the study.

We also enrolled a group of 12 adults (6 female, mean age: 27.2 years; s.d.: 2.2 years). We replaced one adult who exhibited unusual difficulty in handling the experimental setup and exhibited accuracy outside 2 s.d. around the mean of the adult sample in some experimental tasks.

The study was approved by the local ethics committee of Bielefeld University. Adult participants and children’s parents gave written informed consent prior to the experiment. Children gave oral consent after they had received all information prior to the study. Children received a small gift of about 10 € value for their participation. Participants of the adult group received course credit.

### 2.2. Experimental setup, tasks, and procedure

#### 2.2.1. Technical setup

All tasks were implemented in the KINARM End-Point-Lab (Fig. 2A, BKIN Technologies Ltd, Kingston, Ontario, Canada), a robot that serves as a hand manipulandum for hand movements in a 2D horizontal plane in front of the user. Participants grab one or two handles, the position of which is continually recorded at a sampling rate of 1000 Hz. Mobility of the handles can be constrained by the robot, allowing us to restrict movement to the circular trajectories required for our tasks. A monitor projects from above onto a mirror placed above the hands, so that visual information displayed on the mirror appears to be located below the mirror, in the same plane as the hands. We provide videos of the screen display for all tasks in the Supplementary Information (https://osf.io/k2qs7). Depending on their body height, participants were seated on a height-adjustable chair, or stood on an adjustable platform, so that their elbows were angled at 90°. Most children aged 8 and older were tall enough to be seated. We did not observe any difference in children who stood or sat, for instance regarding concentration, motivation, fussiness etc. Independent of whether children stood or sat, they did not complain about fatigue. We took regular breaks to allow recuperation between trials and tasks. For age groups in which there were both standing and seated children, we screened performance data for obvious differences between these two groups but did not detect any; this is, in fact, in keeping with studies that reported only minimal effects of posture on performance when using the Kinarm; Lowrey et al., 2017; Mang et al., 2019). With this arrangement, all participants could reach sufficiently far into the workspace to comfortably move the handles and their arms could move without obstruction. An opaque apron attached to the participant’s neck prevented vision of the arms, and the display mirror was opaque, so that participants could not see their hands. This setup allowed us to fully control visual feedback about hand position via cursors on the screen.

**Fig. 2.**
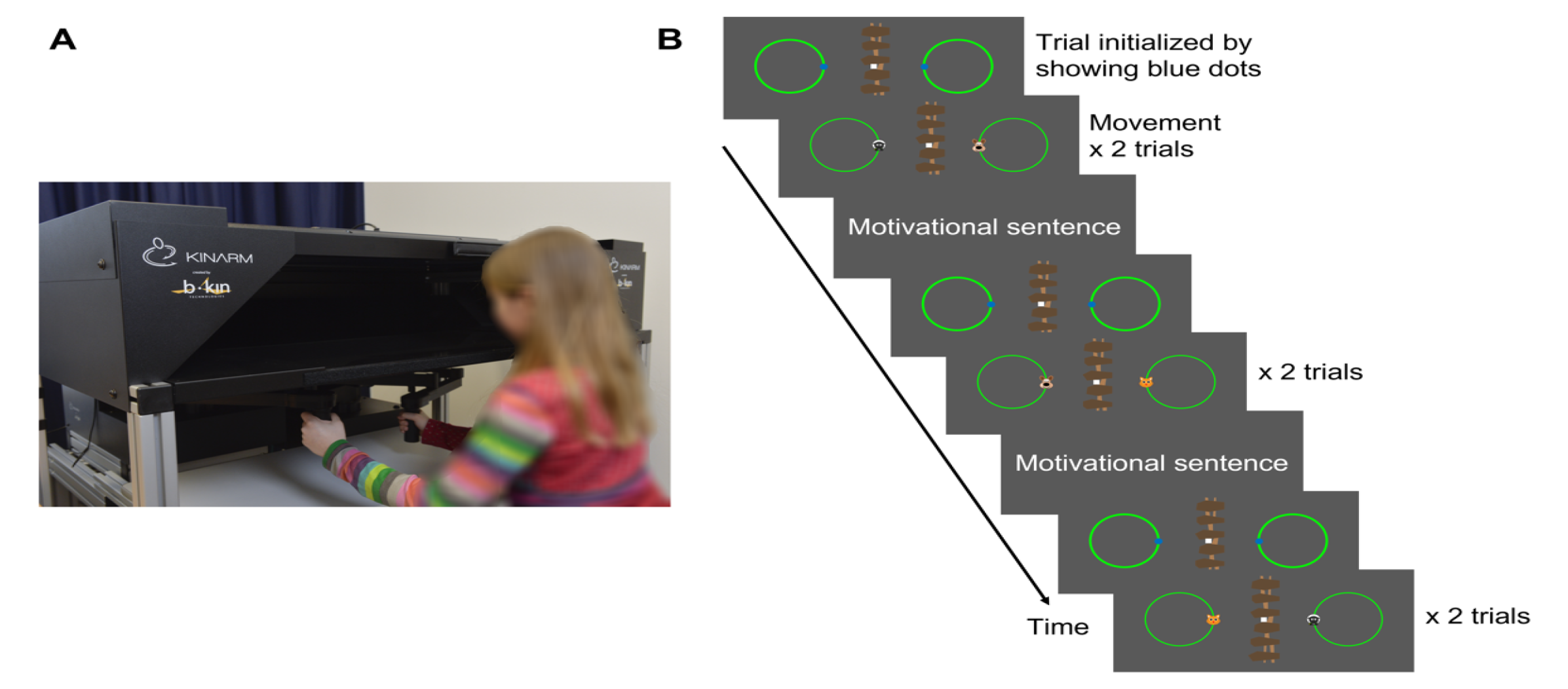
Illustration of the general setup. **(A)** The Kinarm Endpoint Lab device. The display is mounted above the hand-held handles, and visual stimuli appear as if they are located at the level of the hands. An apron (not shown) was attached so that the participant could not see their hands. **(B)** Illustration of the task display. Circles indicated where each hand could be moved. The hands could move freely between trials. Once the participant moved to the blue starting positions, movement was restricted along the circles and participants fixated the central, white square. A short motivational sentence was displayed after every two trials.

#### 2.2.2. Task display

Fig. 2B illustrates the task display. In all tasks, the display showed two same-colored circles of 10 cm diameter. They were 20 cm apart, visually separated by a fence-styled line down the middle of the display. A fixation square was placed in the middle of the fence, at the height of the circles’ midpoints. Some of the 6 tasks required movement of one, and other tasks movement of both hands; whenever the right hand had to be moved, it could only move along the trajectory underneath the right circle outline, and equivalently for the left hand and circle. The starting point for hand movement in a given trial was always the innermost circle position, which were marked with blue, 1 cm diameter dots on the circles. We created child-friendly designs and story lines to support children’s motivation to complete the study. Each child chose one of several stimulus themes that each featured four 2 cm diameter characters to represent hand position (see Fig. 2B & Video in the Supplementary Material with the visual display for each task https://osf.io/k2qs7).

#### 2.2.3. Tasks

We report on 6 tasks in which participants had to move one or both hands in circles. The task setup mimicked turning handles attached to wheels by allowing participants to move the two Kinarm handles along fixed circular trajectories. We present the tasks in the order used for the logic of the paper; children performed the tasks in a different order, which we report below (see Procedure). Tasks 1 and 2 tested unimanual coordination; tasks 3 and 4 assessed bimanual, symmetric coordination; and tasks 5 and 6 addressed sensory selectivity in bimanual motor coordination.

##### Task 1 – Match the Cursor: Unimanual Visual Coordination (Uni-Vis)

Participants had to coordinate a hand with an externally controlled visual signal. They held a Kinarm handle with their dominant hand and rested the non-dominant hand on their lap. The screen was divided in the middle, and there was an area both on the left and right in which circling cursors were presented and circular Kinarm handle movements were possible (see Fig. 2B). Children saw the computer-controlled cursor on the side of the non-dominant hand moving along its circular trajectory. The cursor’s circling frequency was 0.8 Hz, that is, 5 rad/sec or 0.25 m/s. The hand’s position, too, was indicated by a cursor and participants had to match the two cursors symmetrically. We had piloted cursor velocity with young children. There were six trials of 20 s duration each.

##### Task 2: Match the Other Hand: Unimanual Proprioceptive Coordination (Uni-Proprio)

Participants grabbed both Kinarm handles. The Kinarm robot passively moved the non-dominant hand at the same speed as in Task 1. Participants were instructed to relax the passively guided arm and let the robot lead them without actively supporting the movement. Participants had to symmetrically match the passive movement of their non-dominant hand with their dominant hand. No visual feedback of either hand’s position was available. There were six trials of 20 s duration each.

##### Task 3 – Bimanual 1:1 Coordination Without Vision (Bi-NoVis)

Participants had to move the two unseen hands symmetrically in six trials of 10 s duration each. There were no cursors for the hands and so the task appeared visually identical to Task 2 (Uni-Proprio).

##### Task 4 – Bimanual 1:1 Coordination With Vision (Bi-Vis)

Participants had to move the two hands symmetrically while receiving feedback about hand position by cursors on the screen so that he task appeared visually identical to Task 1 (Uni-Vis). They performed six trials of 10 s duration each. Piloting had revealed that Tasks 3 and 4 were quite easy to perform and we chose a short trial duration to avoid children getting bored and demotivated.

##### Task 5 – 2:3 Coordination With Veridical Hand Position Feedback (2:3-NoTransform)

Participants had to concurrently perform three circles with their dominant, and two with their non-dominant hand. Hand position was indicated by visual cursors, as in Uni-Vis and Bi-Vis. Accordingly, if the task was performed correctly, the two cursors moved asymmetrically. This kind of task is very difficult – often impossible – even for adults and constituted a baseline 2:3 coordination task. Participants performed six trials of 20 s duration each.

##### Task 6 – 2:3 Coordination With Transformed Feedback (2:3-Transform)

Participants had to perform 2:3 coordination as in Task 5 (2:3-NoTransform). However, cursor position did not reflect the hands’ veridical location but was transformed to appear symmetrical: For the dominant hand, the cursor completed one full circle on the display while participants performed 3 circles with their hand. For the non-dominant hand, the cursor completed one circle while the hand circled twice. Accordingly, participants could correctly perform the task by maintaining symmetrical visual feedback on the display. We explained this task feature in detail to participants. They performed 3 x 6 trials of 20 s duration each.

We conducted more trials in this task, because we expected that participants would learn the required asymmetrical coordination with some practice, based on previous reports (Chiou & Chang, 2016; Kovacs et al., 2009, 2010; Kovacs & Shea, 2011). We tested potential learning in a seventh task, Retention, in which 2:3 transformed visual feedback was provided only at the beginning of each trial and then removed. With such a procedure, a previous study reported coordination learning in adults (Kovacs & Shea, 2011). However, in our study here, neither children nor adults were able to perform this seventh task correctly. This finding indicates that none of our participants learned the 2:3 coordination independent of visual feedback. We omit a detailed report of this seventh task for the sake of brevity (but see supplementary information https://osf.io/grnbx).

###### Trial structure

For all tasks, each trial began with the participant moving their hand(s) to the starting position(s) (Fig. 2B, blue dots). A beep of 150ms duration instructed to begin circling in the inward direction (i.e. left hand clockwise and right hand counter-clockwise) until a beep of 300ms duration signaled the trial’s end. Participants were to look at the fixation square throughout the trial. The experimenter monitored fixation visually and additionally reminded the participant about fixation every two trials. We repeated trials if the participant did not fixate as instructed. For all tasks, participants performed three practice trials during which we gave oral feedback on whether task performance complied with the instructions. If necessary, we added up to two additional practice trials. During experimental trials, we gave no feedback about performance. However, if the participant moved in the wrong direction (e.g. counter-clockwise for the left hand), we explained the mistake and repeated the trial. This happened seldom, though more with younger than older children, and was not specific to one task.

After every two trials, we displayed short sentences for 5 s to frame the experiment as a story and support children’s motivation. We read the sentences aloud for children who could not yet read. Sentences playfully framed the tasks and commended them for their continuous effort. Examples of the framing story and sentences are available in the supplementary information (https://osf.io/grnbx).

###### Procedure

Participants first played a short game to acquaint themselves with the Kinarm. Then, they performed the experimental tasks. Participants always started with the bimanual coordination tasks, Bi-NoVis and Bi-Vis (Task 3 and 4 in this report). We used this order because we wanted to avoid that performance without vision would benefit from experience of the visual task if it were done first. Then, Uni-Vis, Uni-Proprio (Task 1 and 2), and 2:3- NoTransform (Task 5) were conducted in randomized order by participants drawing pieces of paper in an envelope. The session ended with 2:3 tasks: 2:3-Transform (Task 6) had to be conducted after 2:3-NoTransform because it potentially involved learning. In turn, Retention (omitted from this report, as explained above) tested learning of 2:3-Transform and, thus, had to be assessed last.

All tasks were 3-5 min long, except for 2:3-Transform, which lasted 12 min. Participants rested between tasks; in the 12 min 2:3 Transform task, they took an additional break in the middle.

Some participants did not perform all tasks. Table 1 lists the number of individuals per year of age and task. We lost some data due to equipment failure; this was especially prevalent for the passive-proprioceptive task, Task 2 (Uni-Proprio), which we could not run for several months; thus, the low number of participants is *not* due to younger children being unable to perform the task, but to the restraint of having to acquire our data within the time range defined by project funding. We decided to nonetheless include the incomplete data of Task 2 because the remaining N is still about 50, and data are complete for some age groups, allowing qualified analysis at least for them.

**Table 1.** Participant numbers for all tasks. Low participant numbers in Task 2 were due to temporary equipment failure that selectively affected this task. Low participant numbers in Tasks 5-6 were due to task difficulty for the younger children.

| Age Group<br>Number of included participants | 4 yo<br>n = 14 | 5 yo<br>n = 12 | 6 yo<br>n = 12 | 7 yo<br>n = 12 | 8 yo<br>n = 12 | 9 yo<br>n = 16 | 10 yo<br>n = 18 | 11 yo<br>n = 12 | 12 yo<br>n = 12 | Adults<br>n = 12 |
| --- | --- | --- | --- | --- | --- | --- | --- | --- | --- | --- |
| **Task 1 – Match the Cursor: Unimanual Visual Coordination – Uni-Vis** | 14 | 12 | 12 | 12 | 11 | 16 | 18 | 12 | 12 | 12 |
| **Task 2 – Match the Other Hand: Unimanual Proprioceptive Coordination – Uni-Proprio** | 1 | 2 | 3 | 5 | 4 | 7 | 6 | 12 | 12 | 12 |
| **Task 3 – Bimanual 1:1 Coordination Without Vision – Bi-NoVis** | 14 | 12 | 12 | 12 | 12 | 16 | 18 | 12 | 12 | 12 |
| **Task 4 – Bimanual 1:1 Coordination With Vision – Bi-Vis** | 14 | 12 | 12 | 12 | 12 | 16 | 18 | 12 | 12 | 12 |
| **Task 5 – 2:3 Coordination With Veridical Hand Position Feedback – 2:3-NoTransform** | 6 | 3 | 11 | 12 | 11 | 16 | 18 | 12 | 12 | 12 |
| **Task 6 – 2:3 Coordination With Transformed Feedback – 2:3-Transform** | 5 | 3 | 11 | 12 | 12 | 16 | 18 | 12 | 12 | 12 |

There are also fewer young participants in the 2:3 tasks (Tasks 5 and 6). In contrast to Task 2, these missing data are due to task difficulty, not to technical problems. Most 4- and 5-year-olds failed in the 2:3 tasks, and some found these tasks very discouraging, leading to low motivation to complete them. Sometime into data acquisition, therefore, we stopped acquiring the 2:3 tasks for children younger than 6 years.

### 2.3. Data processing

We analyzed data in Matlab (version: R2015aSP1). Each hand’s position time series was filtered using a low pass Butterworth filter with a cut-off frequency of 10 Hz.

We base our report on the unsigned phase difference between the two positions that are coordinated: the two hands in bimanual tasks; the visual cursor or the passively guided hand and the active hand in unimanual tasks, and the two transformed locations in the 2:3 tasks. By discounting the sign, we ignore which position leads the other – for instance, whether the dominant hand is ahead of the non-dominant hand or vice versa. Some previous studies have, instead, reported the percentage of performance within a certain interval, for instance +/-30° phase angle as the “percentage of adequate coordination” (Mechsner et al., 2001, 2007) or “time on task“ (e.g. Bingham et al., 2018). Qualitatively, this measure renders similar results as the phase difference (Brandes et al., 2017); we report it in the Supplementary Information (https://osf.io/grnbx) to aid comparison with earlier work.

In theory, analysis of signed phase difference would allow determining whether coordination is based on sensory feedback or its prediction. As processing sensory feedback takes time, the coordinating hand would lag behind the guiding signal. In contrast, when coordination reflects prediction of the guiding signal, the coordinating hand could be ahead of the guiding signal. Results of signed measures were inconclusive and so we report them only in in the Supplementary Information (https://osf.io/grnbx).

We coded hand position as the angle between the starting position and the location along movement direction (see Fig. 3B). Thus, when the hands were at the lowest position, their phase angle was 90°, at the outermost position it was 180° etc. (see Fig. 3A). In the bimanual tasks, we computed the phase angle difference (see Fig. 3C) of the two hands. In the unimanual tasks, we computed the phase angle difference between the actively moved hand and the position of the leading signal, that is, the visual cursor (Uni-Vis) or the passively moved hand (Uni-Proprio). In the 2:3 tasks, we divided the faster hand’s phase angle by 3, the slower hand’s phase angle by 2, and subtracted the two to assess the phase difference relative to the instructed, non-symmetrical movement.

**Fig. 3.**
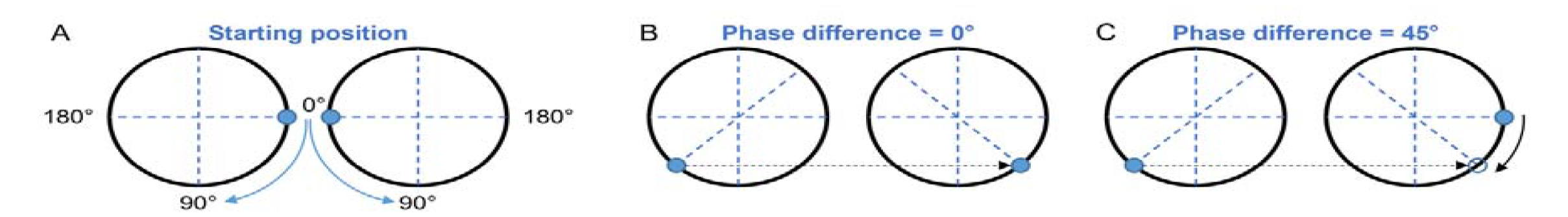
Diagram illustrating phase angle difference calculation. Phase angle was coded along the respective hand’s movement direction, which was mirrored between the two hands. We obtained the phase angle difference by subtracting the dominant hand’s from the non-dominant hand’s phase angle.

Participants performed several circles in every trial. We calculated the average phase difference for each full circle of the dominant hand (see Fig. 4), resulting in several phase difference values per trial (e.g. about 14 circles for each trial in unimanual Tasks 1 and 2). This procedure removes potential artifactual differences between tasks: had we averaged across entire trials, data of longer trials would be smoother due to the larger amount of data. Averaging per performed circle, rather than entering each 1 ms sample into analysis reduced computation time for our statistical models while still retaining sensible and interpretable values for each trial.

**Fig. 4.**
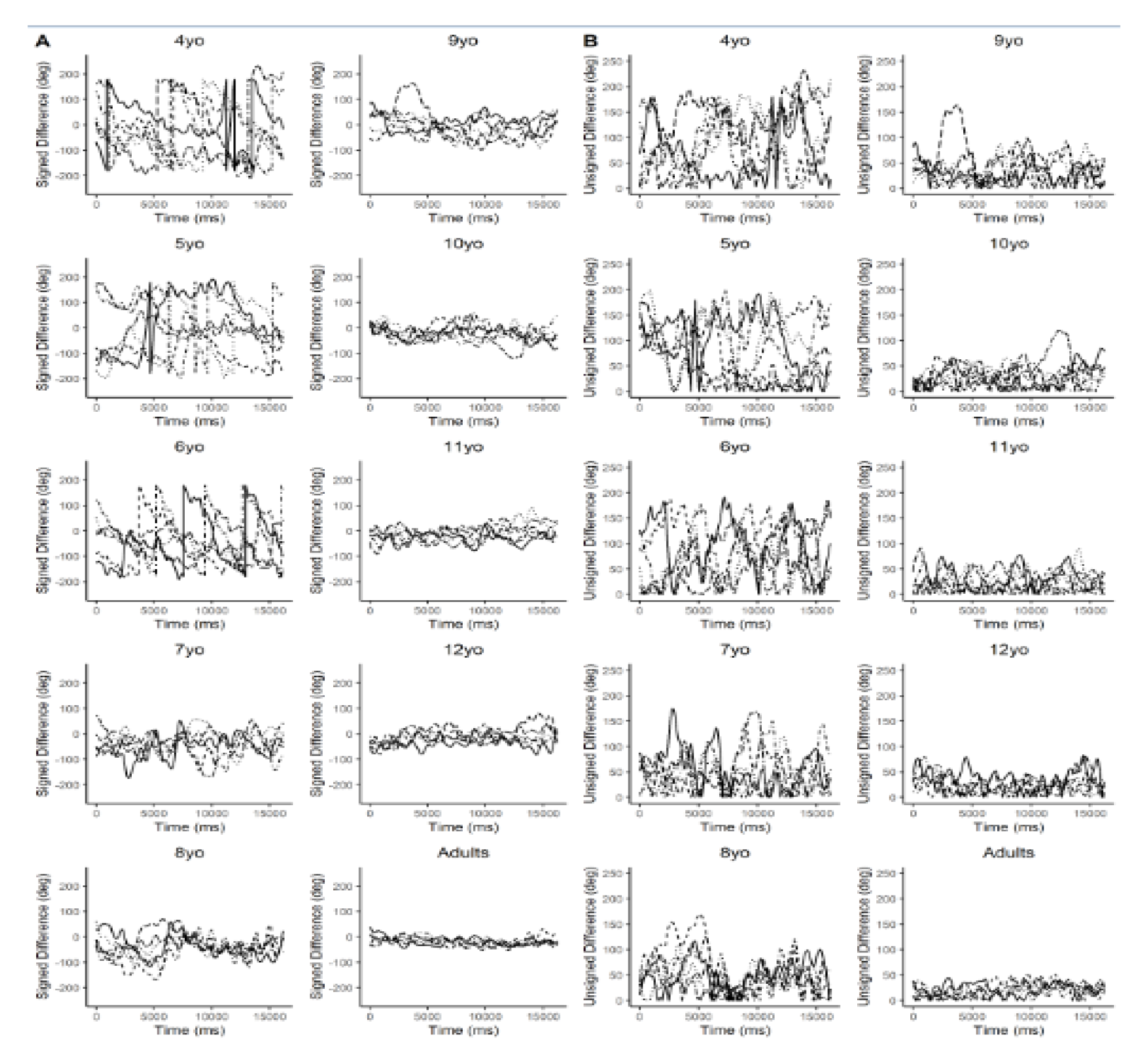
Examples of raw data across entire trials. Time series illustrate performance of a representative participant for each age group for **(A)** the signed phase difference (see supplementary information for the analysis) and **(B)** the unsigned phase difference, in Task 1 (Uni-Vis). This task exhibited the largest performance changes among the 6 tasks across development. All acquired raw data are freely available (https://osf.io/x9u27/). Note that jumps in the signed phased difference observable in the younger groups in A indicate that participants paused one hand while still moving the other one, or were much faster with one hand.

We discarded all data recorded at the trial start until the hands actually started moving to avoid counting this phase as “perfectly coordinated” due to both hands being still. In the unimanual tasks, Tasks 1 and 2, the guiding signal (computer-controlled cursor or passively moved hand) slowly increased its speed over 3 s to avoid startle, and a breaking ramp of 0.5 s. Therefore, we removed the first 3.3 s and last 0.5 s of each trials’ data in these tasks.

### 2.4. Data analysis and statistics

We analyzed data statistically in R (R Core Team, 2018; R: 3.5.1; R Studio: 1.1.456). We implemented multilevel Bayesian modeling with the R package *brms* (Bürkner, 2017).

We addressed our research questions by separately analyzing the symmetrical tasks 1-4 and the 2:3 tasks 5-6, respectively. Analysis of the symmetrical tasks allows gauging the ability to coordinate a hand with visual and proprioceptive information, as well as exploiting motor information of one hand to guide the other. Thus, this analysis addresses the first three aspects discussed in the introduction (Fig. 1A, blue and red shapes; Fig 1B, aspects 1-3). In contrast, the comparison of tasks 5 and 6 specifically addresses the ability of visual synchrony information to coordinate non-symmetrical bimanual movement, thus addressing the third aspect laid out in the introduction (Fig. 1A, ochre shapes; Fig. 1B, aspects 4a,b).

Moreover, we applied three different statistical approaches. First, we used children’s age in months as a continuous predictor of the unsigned phase angle. *Age* was mean-centered, allowing to interpret potential interactions with reference to the sample’s mean age. We excluded adults in this model, because their high performance plus the large age gap between their and the children’s age in months, might have led to artificial correlations of age with coordination performance. This approach identifies general performance trends across age, but does not afford comparing specific age groups. Small irregularities, such as a temporary decline of performance, say, at ages 7 or 8, as have been previously reported, would be smoothed out by the continuous predictors.

Therefore, in our second approach, we devised separate models in which Age in years was a entered as a categorical variable. Here, we included adults as one group, so that we could compare post-hoc, for each task, whether children at each age performed as successfully as adults. Moreover, it was possible to contrast children’s performance between the different ages, allowing us to detect investigate reversals and performance plateaus which would be missed by a continuous, linear predictor.

Third, we tested whether performance was related across (some of) the six tasks. Given that one expects performance in any task to improve with age, all tasks will be trivially related via age. Therefore, we computed correlations between tasks while partialling out age to isolate any potential age-independent performance relationships across tasks.

We implemented the first and second analyses as Bayesian mixed models. Mixed models cope well with different numbers of observations across conditions, small samples, and groups of different sample size (Boisgontier & Cheval, 2016). All models used the unsigned phase difference, averaged per full circle of the dominant hand, as dependent measure (see above, Section 2.3).

*Model 1* used as predictors the within-subject, categorical variable *Task* (Task 1-4: Uni-Vis, Uni-Proprio, Bi-NoVis and Bi-Vis,), the between-subject, continuous variable *Age* (in months), and the interaction between *Task* and *Age*. The model was defined as: Phase_Difference ∼ Task × Age + (1|subject), thus allowing random effects per subject.

*Model 2* used the predictor *Age* grouped by year as a categorical variable. The model was defined as: Phase_Difference ∼ Task × Age_Group + (1|subject).

*Model 3* was defined just like Model 1, except that Task comprised the 2:3 tasks (Task 5-6: 2:3-NoTransform, 2:3-Transform).

Similarly, *Model 4* was defined just like Model 2, except that Task comprised the 2:3 tasks (Task 5-6: 2:3-NoTransform, 2:3-Transform).

Bayesian mixed models have the additional advantage of allowing to assess null effects. The Highest Density Interval (HDI) represents the 95% interval of a parameter estimate’s posterior distribution, that is, the parameter distribution estimated based on the experimental data. When the HDI clusters around 0, operationalized as the complete HDI falling within +/- 0.1 standard deviations of the dependent variable (termed the Region of Practical Equivalence, ROPE, of effect sizes; Kruschke, 2011, 2018), one concludes that the parameter does not have an effect. On the flip side, if the HDI does not fall within the ROPE, it is considered credible that the parameter has an effect. In some cases, only part of the HDI might fall within the ROPE. This inconclusive result prohibits concluding both a difference between conditions and equivalence. When we applied ROPE analysis to compare children’s to adults’ performance, we interpreted this latter case as indicating that children’s performance did not yet resemble adult performance, given that the two groups’ performance could not be considered equivalent.

For each main effect and interaction, we report its estimate (β) and its HDI_95%_. We used the R package *sjstats* (Lüdecke, 2018) to compute the ROPE for each model and infer the presence of statistical effects (ROPE limits for each model: Model 1: [-4.1; 4.1]; Model 2: [-3.9; 3.9]; Model 3: [-5.4; 5.4]; Model 4: [-5.4; 5.4]; numbers refer to the dependent variable: phase difference in degrees). To get meaningful values for models with a continuous predictor *Age*, we standardized the posterior distribution of this covariate (and respective interactions) by multiplying the Age covariate and its interactions, as well as the respective HDIs, by the standard deviation of subjects’ age; this procedure standardizes the estimate range, similarly as z-standardization. The SD of subjects’ age differed between Tasks 1-4 (Models 1 and 2: 30.3) and Tasks 5-6 (Models 3 and 4: 25.3) because fewer children completed tasks 5-6 than 1-4 (see Methods).

Model fitting used a Gaussian likelihood and the *brms* package’s default weakly-informative, improper, flat priors for population-level (“fixed”) effects. The estimation of parameters’ posterior distribution was obtained by Hamiltonian Monte-Carlo sampling with 4 chains, 1000 sample warmup, and 11000 iterations, and checked visually for convergence (high ESS and Rhat ≈ 1).

We implemented our third analysis, the assessment of whether there were age-independent developmental dependencies across tasks, as Bayesian partial correlations. We correlated the unsigned phase angle between pairs of tasks, but removed the effect of age as implemented in the R package *BayesMed* (Wetzels & Wagenmakers, 2012) using a default Jeffrey’s-Zellner-Siow prior set-up. The obtained Bayes Factor (BF) expresses the odds of the H1, a true correlation, compared to the H0 of a zero correlation. A BF between 3 and 10 indicates moderate evidence for the H1, while values >10 above are considered strong evidence (Wetzels et al., 2011).

## 3. RESULTS

### 3.1. Uni-and bimanual, symmetric coordination

This analysis pertains to Tasks 1-4. The underlying study design and logic are illustrated in Fig. 1A and 1B (circles 1-3).

#### 3.1.1. Model 1: Age as continuous predictor

We assessed the variability of children’s coordination performance in a model that predicted the unsigned phase difference between the two coordinated signals – a visual or proprioceptive signal and a hand (Tasks 1 & 2, Uni-Vis & Uni-Proprio, unimanual), or the two hands (Tasks 3 & 4, Bi-Vis & Bi-NoVis, bimanual). Fig. 5 illustrates performance in the four tasks included in the model.

**Fig. 5.**
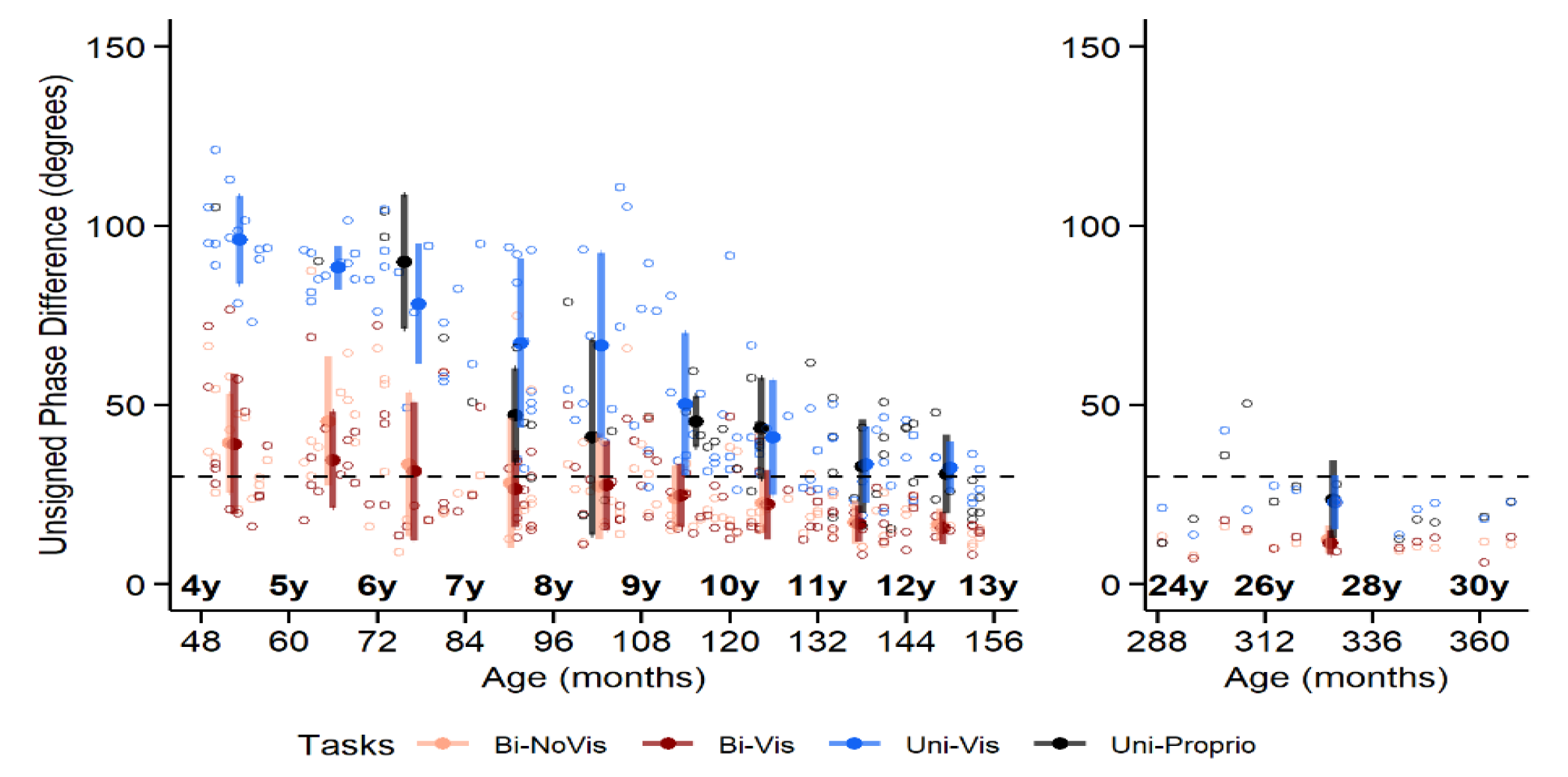
Unsigned phase difference between the two hands in unimanual Tasks 1 and 2 (blue and black) and bimanual Tasks 3 and 4 (dark and light red) for children (left panel) and adults (right panel). The dashed lines represent the 30° threshold under which we considered subjects to perform well (see supplement for analyses using this threshold to define the dependent variable: https://osf.io/grnbx). Colored circles represent the mean for each age group; bars show the s.d. Fewer than half of the 4-8 year-old participants performed Task 2 (Uni-Proprio; black), the means are thus likely to be non-representative (means in Uni-Proprio are not shown for 4- to 5-year-old children, because n ≤ 2).

In the two unimanual tasks, participants had to synchronize the movement of their dominant hand with a circling dot (Uni-Vis) or with their passively moved, non-dominant hand (Uni-Proprio). β estimates reflect improvement with age: Negative slope values indicate improving performance, as the phase difference between hand and sensory stimulus decreased by the number of degrees specified by β_Age_ with each standardized age unit (30.3 months in Model 1, see Methods). Slope coefficients for the two unimanual tasks were indeed negative and different from 0 (Uni-Vis: β_Age_ = −21.7, HDI_95%_ = [-23.7; −19.6], Fig. 5: blue; Uni-Proprio: β_Age_ = −19.4, HDI_95%_ = [-21.6; −16.9], Fig. 5: black; see Table 2), thus indicating that children performed them better with increasing age. There was no credible difference for the contrast between the two tasks, indicating that the Intercept coefficient was similar for Uni-Proprio than Uni-Vis (difference of Uni-Proprio and Uni-Vis, β = −4.0, HDI_95%_ = [-5.5; −2.5]; see Table 2). Moreover, there was no credible *Task* × *Age* interaction contrast either (β = 2.3, HDI_95%_ = 0.8; 3.8]; see Table 2), indicating that the slope did not differ between unimanual tasks. Thus, the two unimanual tasks did not differ in any of their estimated parameters.

**Table 2.**
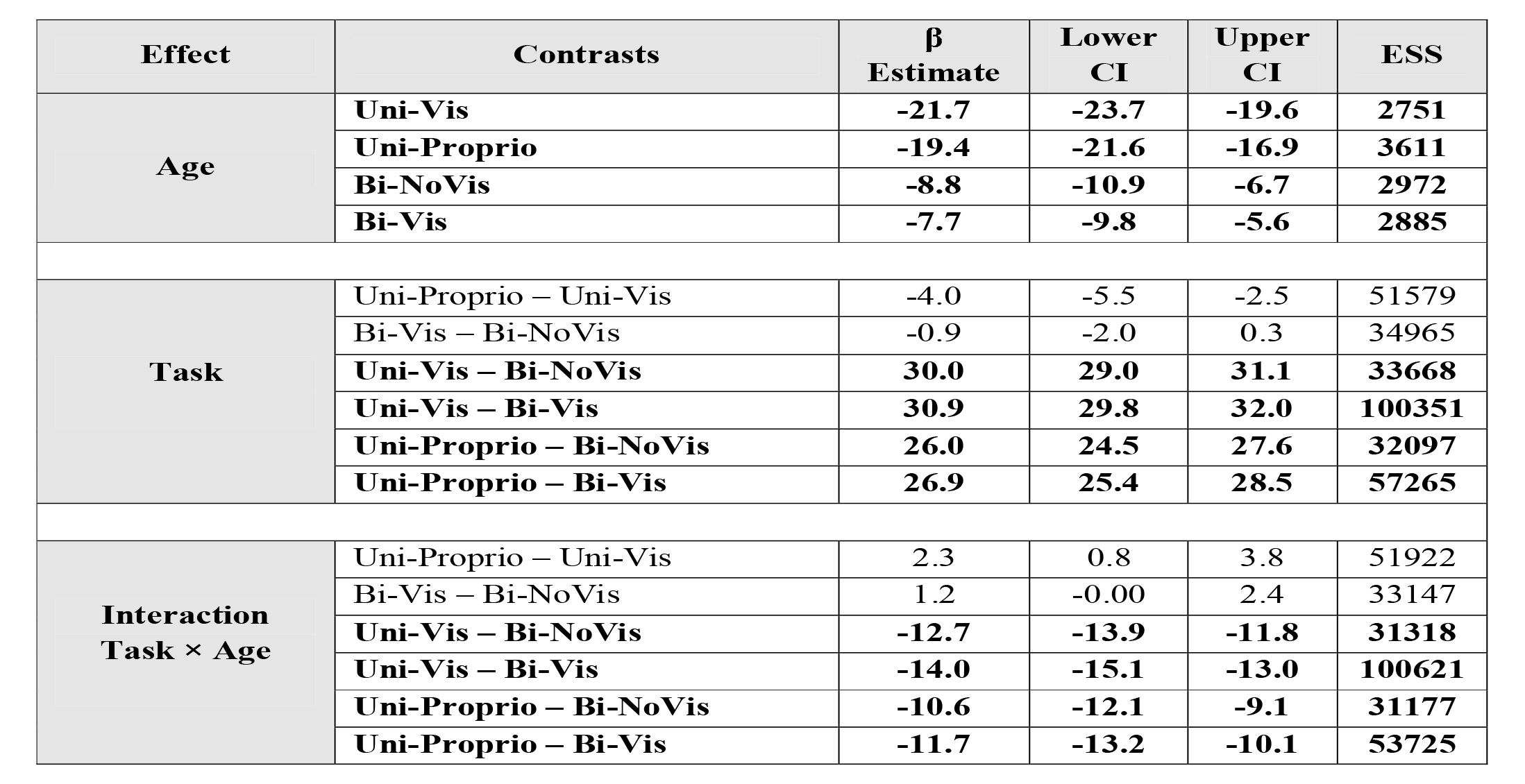
Beta Estimates reflecting the comparison between Tasks 1 to 4 in Model 1. For Age, negative β Estimates indicate that performance was better with higher age (smaller error for each standardized unit of Age of 30.3 months, see Methods). For Task, positive β Estimates indicate that Intercept was higher in the first task of each line, meaning that for this specific task, there was a higher error at younger ages. The interaction Task × Age reflects differences in steepness of improvements with Age. The Lower and Upper Confidence Interval (CI) values constitute the 95% High Density Interval (HDI), which represent the interval in which 95% of the values will be found. Credible differences are indicated in bold (HDI outside of the Region Of Practical Equivalence in which there is no effect [-4.1; 4.1]). ESS stands for the Effective Sample Size as calculated during Bayesian model fitting.

| Effect | Contrasts | $\beta$<br>Estimate | Lower<br>CI | Upper<br>CI | ESS |
| --- | --- | --- | --- | --- | --- |
| Age | Uni-Vis | -21.7 | -23.7 | -19.6 | 2751 |
|  | Uni-Proprio | -19.4 | -21.6 | -16.9 | 3611 |
|  | Bi-NoVis | -8.8 | -10.9 | -6.7 | 2972 |
|  | Bi-Vis | -7.7 | -9.8 | -5.6 | 2885 |
| Task | Uni-Proprio – Uni-Vis | -4.0 | -5.5 | -2.5 | 51579 |
|  | Bi-Vis – Bi-NoVis | -0.9 | -2.0 | 0.3 | 34965 |
|  | Uni-Vis – Bi-NoVis | **30.0** | **29.0** | **31.1** | **33668** |
|  | Uni-Vis – Bi-Vis | **30.9** | **29.8** | **32.0** | **100351** |
|  | Uni-Proprio – Bi-NoVis | **26.0** | **24.5** | **27.6** | **32097** |
|  | Uni-Proprio – Bi-Vis | **26.9** | **25.4** | **28.5** | **57265** |
| Interaction<br>Task × Age | Uni-Proprio – Uni-Vis | 2.3 | 0.8 | 3.8 | 51922 |
|  | Bi-Vis – Bi-NoVis | 1.2 | -0.00 | 2.4 | 33147 |
|  | Uni-Vis – Bi-NoVis | **-12.7** | **-13.9** | **-11.8** | **31318** |
|  | Uni-Vis – Bi-Vis | **-14.0** | **-15.1** | **-13.0** | **100621** |
|  | Uni-Proprio – Bi-NoVis | **-10.6** | **-12.1** | **-9.1** | **31177** |
|  | Uni-Proprio – Bi-Vis | **-11.7** | **-13.2** | **-10.1** | **53725** |

In the two bimanual tasks, participants had to symmetrically rotate their hands, with (Bi-Vis) or without (Bi-NoVis) additional visual feedback of their hand position. Both slope coefficients were negative and different from 0 (Bi-NoVis: β_Age_ = −8.8, HDI_95%_ = [-10.9; −6.7]; Bi-Vis: β_Age_ = −7.7, HDI_95%_ = [-9.8; −5.6]; see Table 2). In equivalence to the unimanual tasks, negative slope values indicate that the phase difference between the two hands decreased by the number of degrees specified by β_Age_ with each standardized age unit of 30.3 months (Fig. 5; light and dark red dots). There was neither a credible difference between the intercepts of the two bimanual tasks, nor a *Task* × *Age* interaction, indicating that the parameters of the two bimanual tasks were not credibly different (see Table 2 for a summary of all model estimates). Thus, performance and its improvement with age were similar in the two bimanual tasks.

The two unimanual tasks exhibited large effects of *Task* and *Task × Age* interactions when contrasted with the bimanual tasks (see Table 2 for all model estimates). Unimanual intercepts were higher, indicating that the performance was worse at younger ages (higher error), and the negative slopes were steeper indicating larger improvement with age, in the unimanual than in the bimanual tasks.

#### 3.1.2. Model 2: Age grouped by year

In the second model, we grouped children into yearly cohorts and contrasted performance between tasks for each cohort as well as for the adult group. Results of this analysis are illustrated in see Fig. 6. Table 3 presents estimates for the four tasks.

**Fig. 6.**
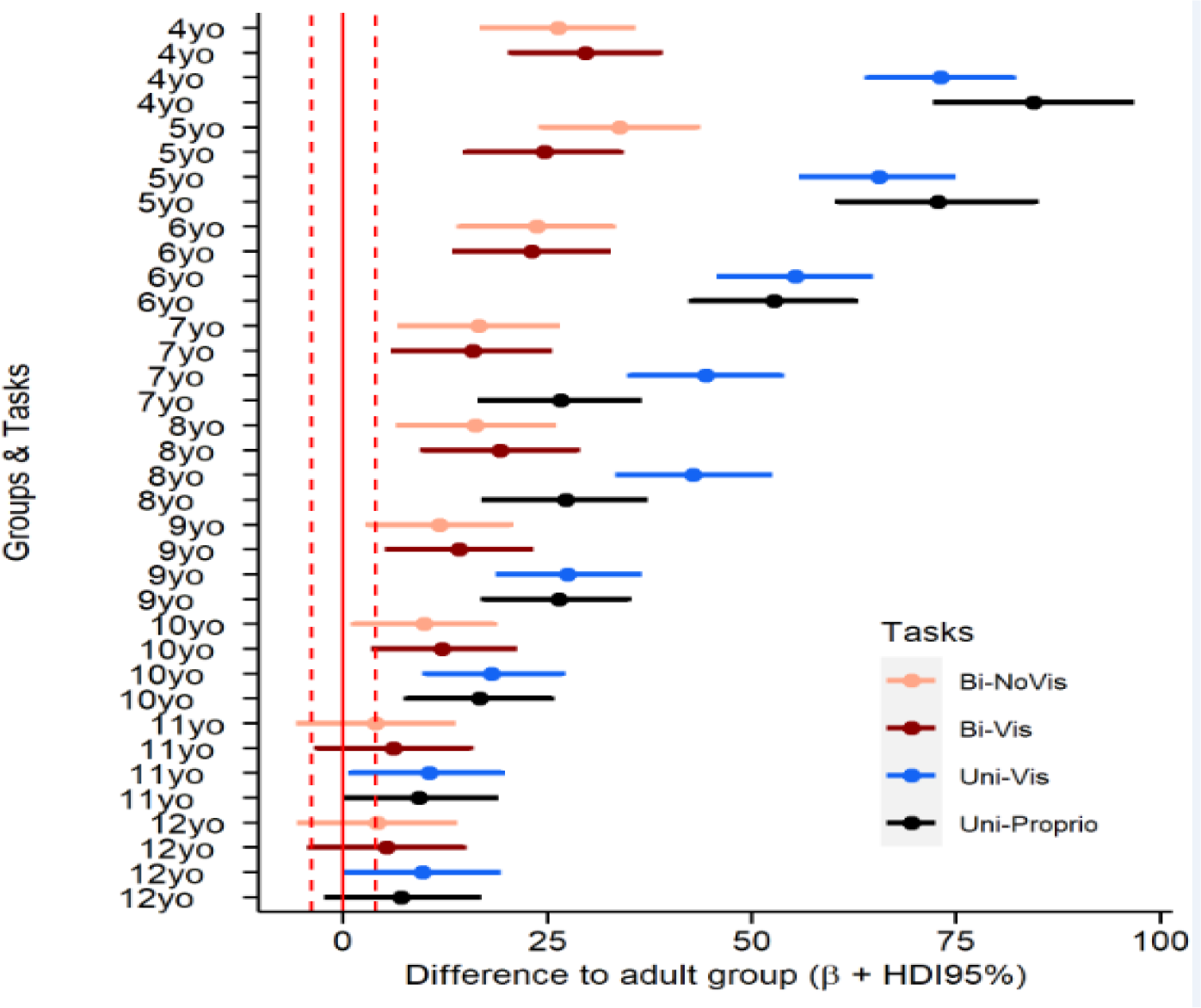
Illustration of age group effects in Tasks 1-4 in Model 2, with Task and Age group as independent variable. The area between whiskers represents the highest density interval (HDI_95%_), that is the area with 95% of the values, and the dot represents the mean β estimate. Following this model, with the adult group as the reference, the differences between adults and the other groups are credible if they do not include the Region Of Practical Equivalence (ROPE: [-3.9; 3.9]; indicated by the vertical dashed lines).

**Table 3.**
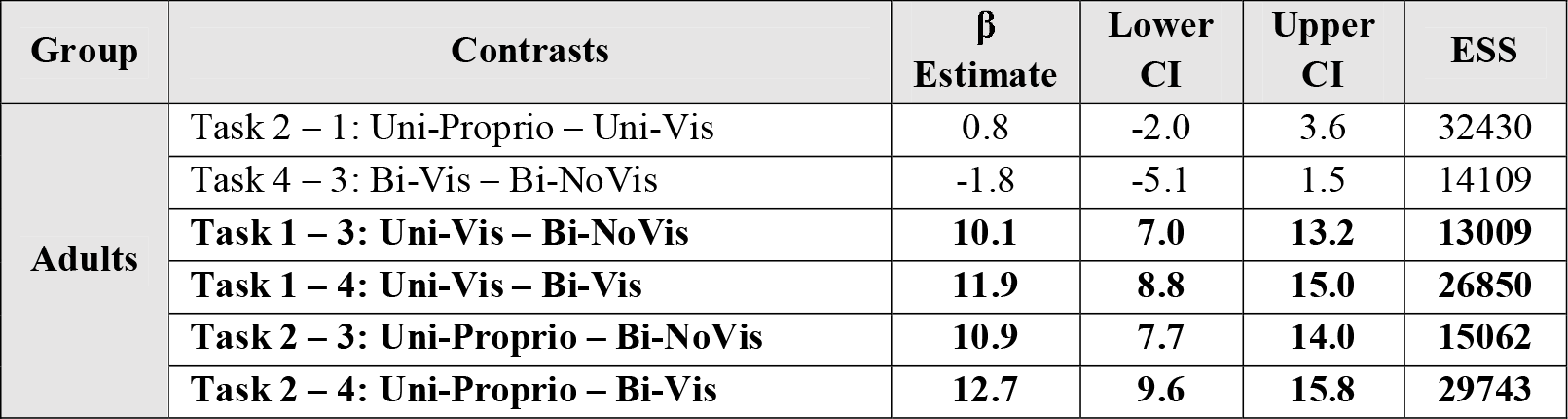
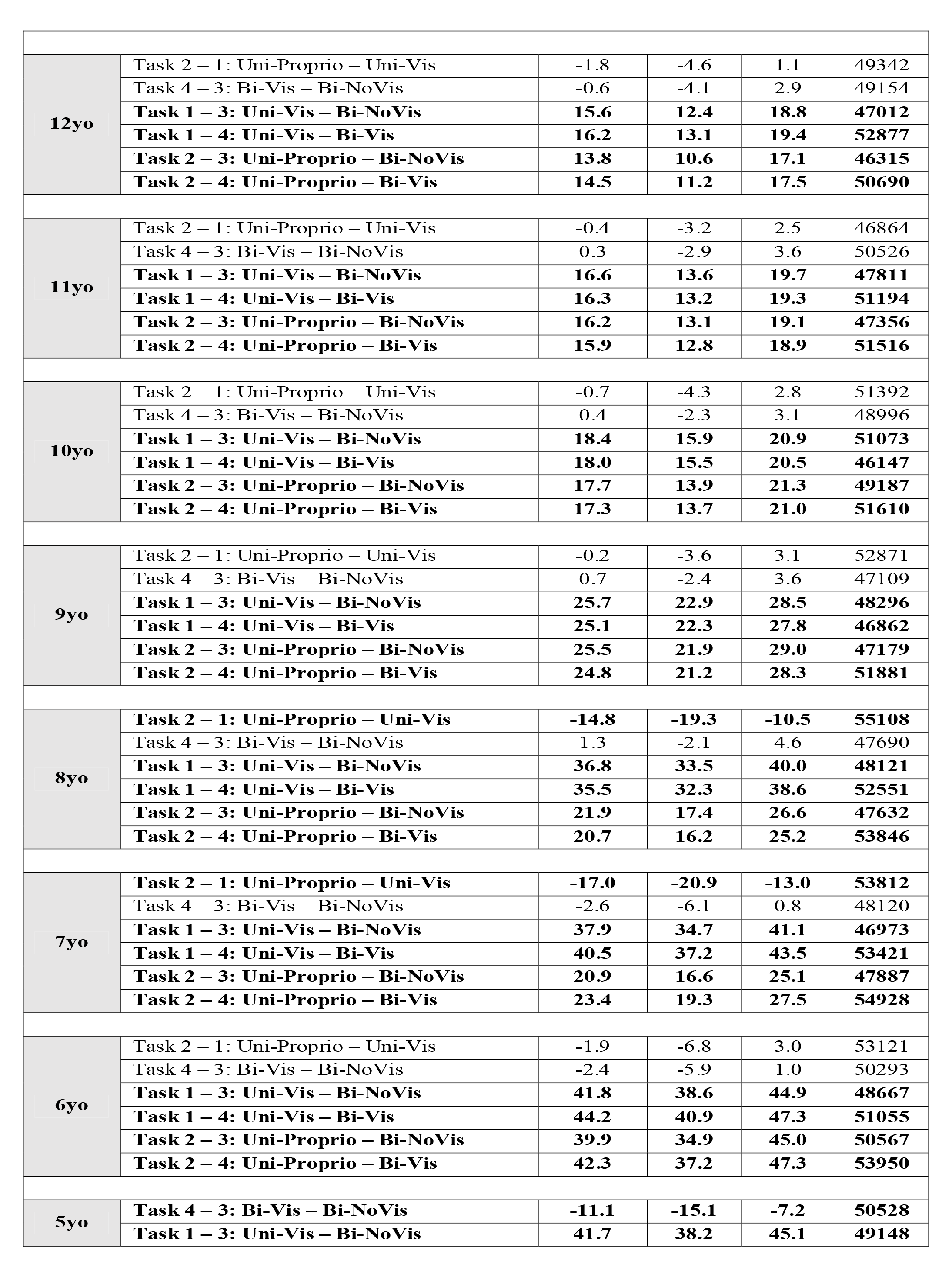
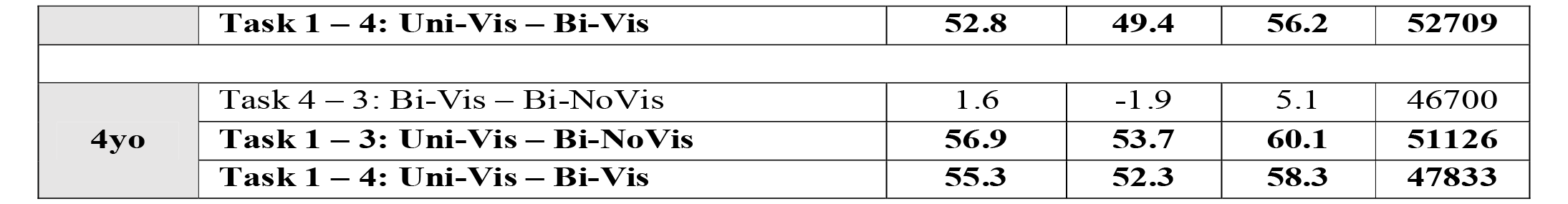
Beta Estimates of the within comparison between Tasks 1 to 4 for each age group in Model 2. Comparison with Task 2 (Uni-Proprio) was not processed for children aged 4-5 years as only few children per group performed the task (≤ 2). Positive β Estimates indicate that performance was worse in the first task of each line. The Lower and Upper Confidence Interval (CI) values constitute the 95% High Density Interval (HDI), which represent the interval in which 95% of the values will be found. Credible differences are indicated in bold (HDI outside of the Region Of Practical Equivalence in which there is no effect [-3.9; 3.9]). ESS stands for the Effective Sample Size as calculated during Bayesian model fitting.

Coordination was better for bimanual than for unimanual tasks across all experimental groups, including adults. This difference was more pronounced for younger than for older children. In the unimanual tasks, 7-year-old (Uni-Proprio – Uni-Vis: β_Task_ = −17.0, HDI_95%_ = [-20.9; - 13.0]; see Table 3) and 8-year-old children (Uni-Proprio – Uni-Vis: β_Task_ = −14.8, HDI_95%_ = [- 19.3; −10.5]; see Table 3) were better in Uni-Proprio than in Uni-Vis (compare black and blue dots in Fig. 3), while there was no difference between the tasks for older participants. Yet, given the low number of children aged 4-7 that participated in Uni-Proprio (due to technical failure, see Methods), we do not think the differences between the two tasks are interpretable.

Performance between the two bimanual tasks did not credibly differ at any age, including adults. There was a single exception for the cohort of age 5 (Bi-Vis – Bi-NoVis: β_Task_ = −11.1, HDI_95%_ = [-15.1; −7.2]; see Table 3), for which coordination was worse when no visual feedback was available (compare light and dark red large dots in Fig. 5).

With this second model, we also assessed at what age children performed comparable to adults (see Table S1 in the supplementary information: https://osf.io/grnbx). For the unimanual tasks, children up to age 10 performed credibly worse than adults. In contrast, there was no credible performance difference for 11 and 12-year-old children compared to adults (see Fig. 6, the blue and black lines cross the red dashed lines only for the 11 and 12- year-old children). However, there was also not sufficient evidence for practical equivalence between 11-12 year-olds and adults, with part of the posterior parameter distribution outside the ROPE. The current data, thus, do not allow us to conclude that development is complete by age 12.

Children performed better in the bimanual than in the unimanual tasks. For Bi-NoVis, children up to age 8 showed credibly higher phase error than adults (see Fig. 6; light and dark red lines cross the vertical dashed lines only for the younger groups). In contrast, for 9 to 12- year-old children, performance neither differed credibly from adults, nor could it be considered practically equivalent. For Bi-Vis, performance of children up to age 9 differed credibly from adults, whereas 10-12 year-olds neither differed from adults, nor did they perform equivalently.

### 3.2. Sensory dominance in bimanual motor coordination

These analyses pertain to Tasks 5 and 6; the underlying study design and logic are illustrated in Fig. 1A and 1B (circles 4a, 4b).

#### 3.2.1. Model 3: Age as continuous predictor

As the third aspect of motor control, we investigated the ability of children to perform asymmetric coordination.

Fig. 7 shows that performance in 2:3-Transform improved with age (green dots), as reflected by a negative, non-zero slope coefficient (β_Age_ = −14.0, HDI_95%_ = [-16.4; −11.4]). The negative slope value indicates that the phase difference between hands decreased by the respective amount of degrees with each standardized age unit of 25.3 months; this unit differed from the previous analyses because most young participants did not perform the 2:3 tasks so that the included participants differed between Models 1/2 and 3/4. In contrast, performance in the 2:3-NoTransform did not credibly vary with age (Fig. 7, yellow dots; β_Age_ = −4.7, HDI_95%_ = [- 7.1; −2.2]). There was a credible difference between tasks for both *Task* and *Task* × *Age* interaction contrasts, indicating that the intercept and rate of improvement was smaller for 2:3-NoTransform than 2:3-Transform (β_Task_ = −13.6, HDI_95%_ = [-14.9; −12.3]; β_Interaction_ = −9.4, HDI_95%_ = [-10.6; −8.1]). During a 2:3 coordination, the correct phase difference between the two hands changes from moment to moment. If a participant incorrectly performed symmetrical (i.e., 1:1) coordination, a pattern that often emerges automatically, then the resulting average error will be about 83°. In Fig. 7, we plot this error as a reference line, together with participants’ performance. Note, that performance in 2:3-NoTransform (Fig. 7, yellow color) mostly scatters around this 83° line across all ages, including adults. Thus, participants across all age groups were unsuccessful when visual feedback simply indicated the limbs’ current position along their respective circles. In contrast, performance improved in the older groups (Fig. 7, green color), starting around age 10, when visual feedback was transformed such that visual symmetry signaled correct motor performance.

**Fig. 7.**
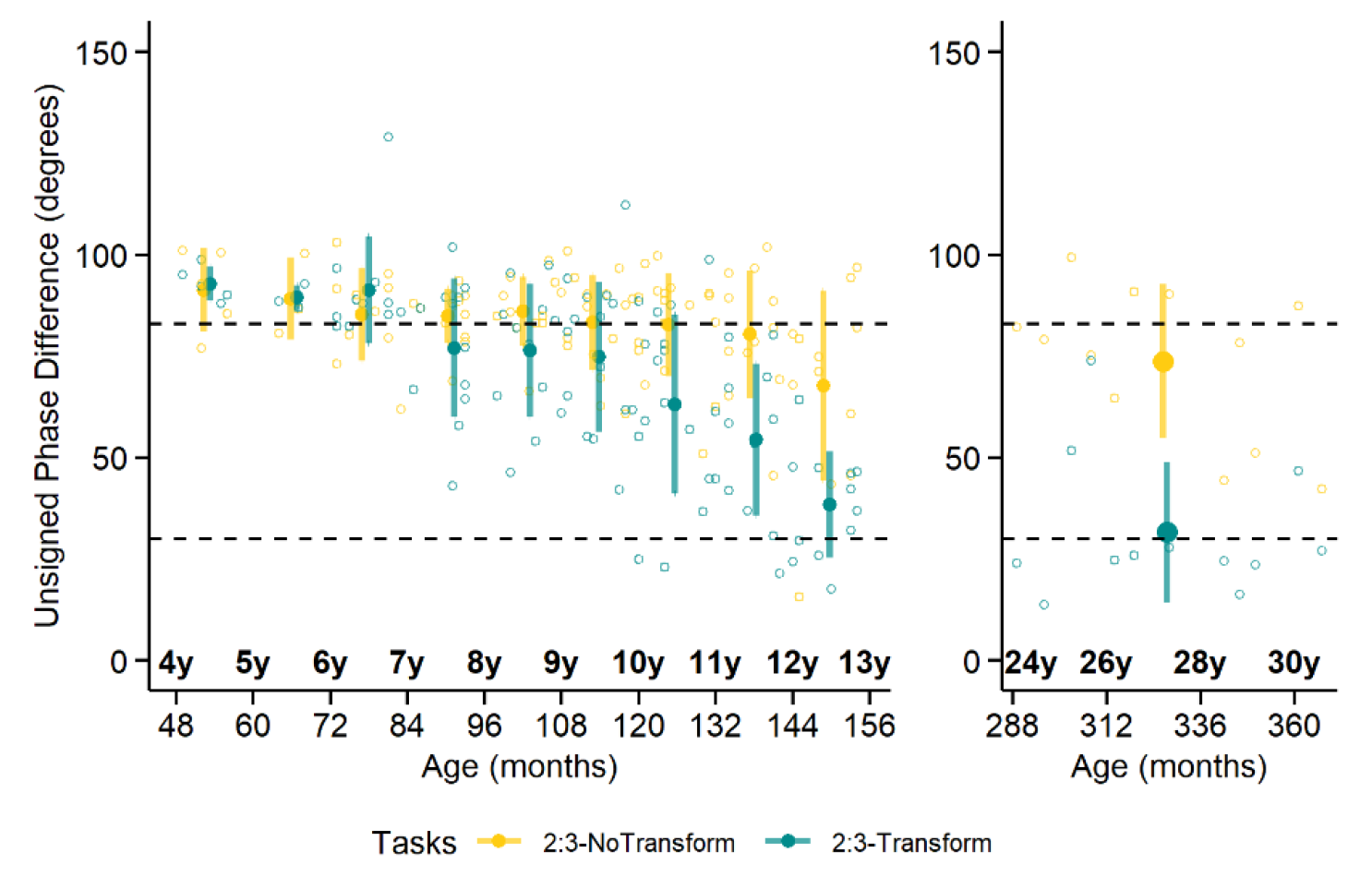
Unsigned phase difference between the two hands in the 2:3 coordination tasks for children (left panel) and adults (right panel). The lowest dashed lines represent the 30° threshold under which we considered subjects to perform well. The higher dashed line (83°) represents the phase angle when participants (incorrectly) perform 1:1 coordination. Coloured circles are the mean for each age group; error bars denote s.d. Fewer than half of the 4-5-year-old participants performed Task 5 (yellow), the means are thus likely to be non-representative.

#### 3.2.2. Model 4: Age grouped by year

In model 4, we contrasted performance between 2:3-NoTransform and 2:3-Transform within each age group. Beta estimates for these comparisons are shown in Table 4. In brief, performance between the two tasks credibly differed in all age groups older than 9, suggesting a consistent use of transformed visual feedback to perform the coordination from this age on. Notably, a credible difference was also present at 7, but not at 8 years. This data pattern may perhaps suggest a temporary performance decrement around age 8, though the numerical differences between ages 7, 8, and 9 are small and, thus, we would not emphasize this result. Children younger than 7 were unable to use transformed feedback (Table 4).

**Table 4.** Beta Estimates of the within comparison between Tasks 5 and 6 for each age group. Negative β Estimates indicate that performance was better in Task 6 (2:3-Transform) than Task 5 (2:3- NoTransform). The Lower and Upper Confidence Interval (CI) values constitute the 95% High Density Interval (HDI), which represent the interval in which 95% of the values will be found. Credible differences are indicated in bold (HDI outside of the Region Of Practical Equivalence in which there is no effect [-5.4; 5.4]). ESS stands for the Effective Sample Size as calculated during Bayesian model fitting.

| Group | $\beta$ Estimate | Lower CI | Upper CI | ESS |
| --- | --- | --- | --- | --- |
| Adults | **-44.6** | **-48.3** | **-40.8** | **9003** |
| 12 yo | **-29.8** | **-34.0** | **-25.9** | **57168** |
| 11 yo | **-27.1** | **-30.7** | **-23.6** | **58690** |
| 10 yo | **-18.3** | **-21.1** | **-15.3** | **58217** |
| 9 yo | **-8.6** | **-11.5** | **-5.7** | **57803** |
| 8 yo | -8.4 | -12.2 | -4.6 | 56434 |
| 7 yo | **-10.6** | **-14.5** | **-6.7** | **59644** |
| 6 yo | 1.5 | -2.3 | 5.4 | 58056 |
| 5 yo | -0.7 | -8.8 | 7.4 | 49878 |
| 4 yo | 1.7 | -4.8 | 8.0 | 43938 |

Comparison of each age group’s performance in 2:3-NoTransform with that of adults did not reveal a credible difference for any age cohort, in line with the above result that 2:3- NoTransform did not improve with age (see Fig. 8 how the yellow lines always cross the vertical dashed lines; statistics reported in Table S2 of the supplementary information: https://osf.io/grnbx). In the 2:3-Transform, the adult group performed better than all age cohorts except the 12-year-olds (see Fig. 8; only the green line of the 12-year-old group crosses the vertical dashed line; β = 9.3, HDI_95%_ = [-1.02; 19.5], but 12-year-olds’ performance neither differed credibly from, nor was practically equivalent to, adult performance.

**Fig. 8.**
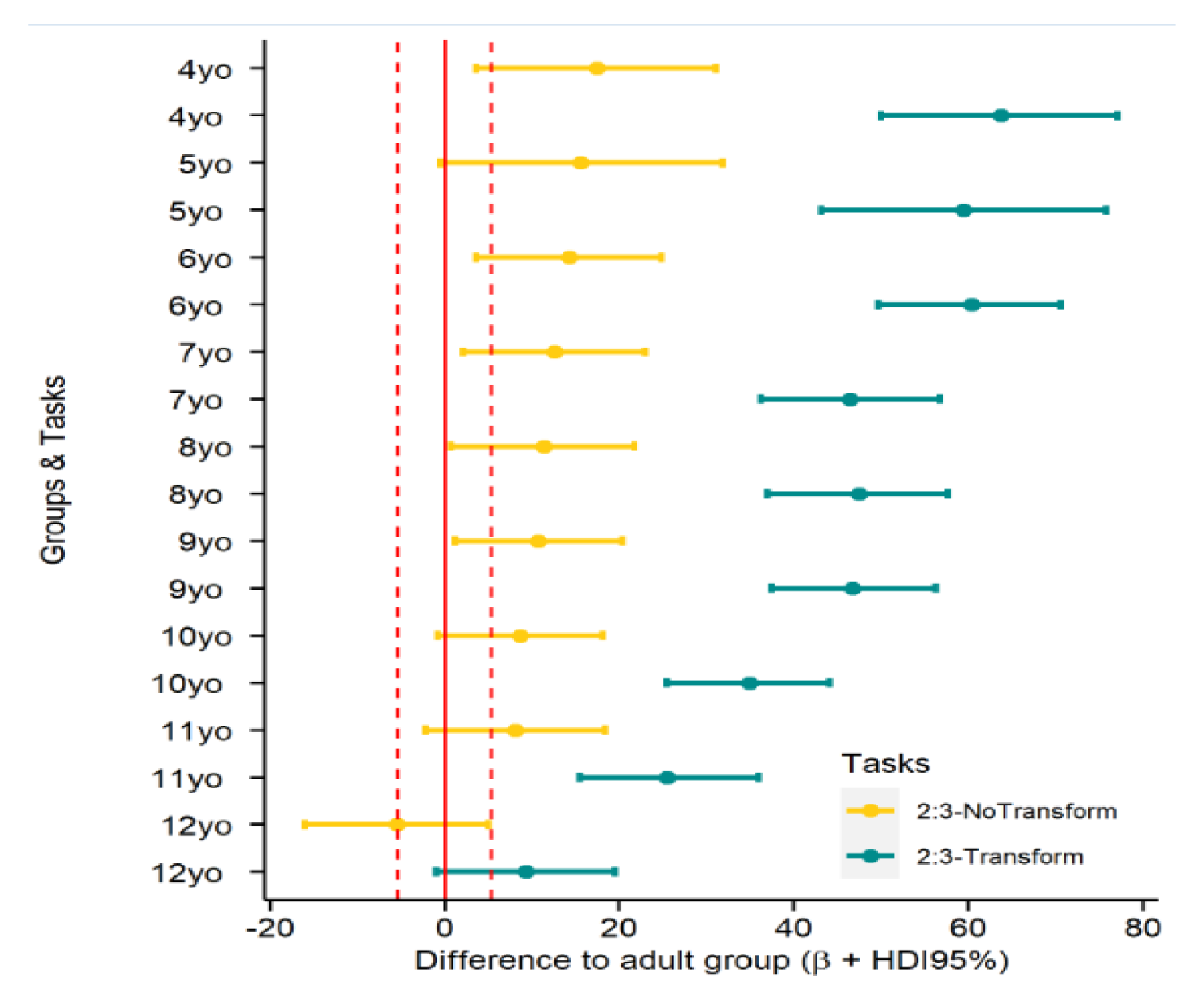
Illustration of age effects in Tasks 5 (2:3-NoTransform) and 6 (2:3-Transform). The area between whiskers represents the highest density interval (HDI95%), that is the area with 95% of the values, and the dot represents the mean β estimate. Following this model, with the adult group as the reference, the differences between adults and the other groups are credible if they do not include the Region Of Practical Equivalence (ROPE: [-5.4; 5.4]; indicated by the vertical dashed lines). Adults performance was not credibly different from any of the groups in the 2:3-NoTransform as all the HDI_95%_ cross the red dashed line. Adults performance was not credibly different from 12-year-old children in 2:3-Transform but was better as compared to all the other groups of children.

### 3.3. Relationship between tasks beyond improvement with age

#### 3.3.1. Relationship between bimanual and unimanual tasks

To assess whether performance in the bimanual and unimanual tasks is related beyond a global age improvement, we computed their correlation with age partialed out (see Table 5). The two unimanual tasks exhibited high correlation (ρ = 0.6; HDI_95%_= [0.3; 0.85]; BF = 264), though the confidence interval was wide due to the low N available for Uni-Proprio. In contrast, correlations between uni-and bimanual tasks were smaller, ranging from .28 to .42, with only the higher correlations exhibiting BFs that suggest reliable relationships. Counterintuitively, higher correlations were obtained for Bi-NoVis with Uni-Vis and for Bi-Vis with Uni-Proprio. For the two visual uni-and bimanual tasks (Bi-Vis and Uni-Vis) and the two proprioceptive tasks (Bi-NoVis and Uni-Proprio), correlations were estimated to be lower. We reiterate that results of analyses that include Uni-Proprio should be regarded with caution due to the low N at younger ages. The lower correlations for this task may be related to this methodological limitation rather than a true difference.

**Table 5.** Two-sided correlations between Tasks 1 to 6, with Age partialed out. Credible correlations are in bold, as indicated by a Bayesian Factor ≥ 3. BF represents the relative evidence of one hypothesis (the presence of a correlation) over another (no correlation). BF between 3 and 10 indicates moderate evidence for a correlation, while everything above indicates strong to extreme evidence (BF>100), and everything below can be considered as anecdotal, suggesting an absence of correlation. Lower and Upper Confidence Interval values constitute the 95% HDI.

| Tasks | Coefficient | Lower CI | Upper CI | BF | Sample (N) |
| --- | --- | --- | --- | --- | --- |
| Uni-Vis / Uni-Proprio | **0.58** | **0.30** | **0.85** | **264** | **51** |
| Bi-NoVis / Bi-Vis | **0.70** | **0.57** | **0.84** | **2.60e15** | **120** |
| Bi-NoVis / Uni-Vis | **0.42** | **0.18** | **0.67** | **16.5** | **119** |
| Bi-NoVis / Uni-Proprio | 0.29 | -0.06 | 0.64 | 0.44 | 51 |
| Bi-Vis / Uni-Vis | 0.28 | 0.02 | 0.54 | 0.60 | 119 |
| Bi-Vis / Uni-Proprio | **0.40** | **0.10** | **0.69** | **3.4** | **51** |
| 2:3-NoTransform / 2:3-Transform | **0.40** | **0.17** | **0.64** | **23.2** | **100** |
| Uni-Vis / 2:3-Transform | **0.30** | **0.08** | **0.52** | **3.0** | **100** |
| Uni-Proprio / 2:3-Transform | 0.33 | 0.04 | 0.62 | 1.49 | 50 |
| Bi-NoVis / 2:3-Transform | 0.08 | -0.09 | 0.26 | 0.17 | 101 |
| Bi-Vis / 2:3-Transform | 0.10 | 0.07 | 0.27 | 0.22 | 101 |

Note, for Uni-Proprio, participants had to let their non-dominant hand be passively guided. It is theoretically possible that children did not conform to this instruction. If children had, instead, actively moved their non-dominant hand, then they would effectively have performed a Bi-NoVis task. Both the much worse performance of the unimanual tasks, and the lower correlations between unimanual and bimanual tasks than those between the bimanual tasks, indicate that children did not perform bimanual coordination and, thus, did not actively synchronize their hands, during Uni-Proprio.

#### 3.3.2. Relationship between bimanual tasks

In the bimanual tasks, the partial correlation was 0.70 (HDI_95%_ = [0.57; 0.84]; n =120; two-sided test; BF = 2.60e15; Table 5). This high correlation implies that a child that was good in one task was also good in the other, and, thus, suggests that the two tasks are driven by common underlying mechanisms.

#### 3.3.3. Relationship between asymmetric coordination tasks

Finally, we asked how performance in the 2:3 tasks was related amongst each other and with the 4 symmetrical tasks within participants (see Table 5). Correlation between the two 2:3 tasks with age partialed out revealed a credible correlation of 0.40 (HDI_95%_ = [0.17; 0.64]; two-sided test; n = 100, BF = 23.2). Thus, a child who performed well with veridical feedback performed well also with transformed feedback, and a child who performed badly in the veridical feedback task gained less from transformed feedback. Correlations with the symmetrical tasks were overall low. They were lowest with the bimanual tasks, and higher, though moderate in size, for the unimanual tasks.

## 4. DISCUSSION

Our study investigated four aspects of motor control in children aged 4-12 through 6 uni-and bimanual circling tasks acquired in a cross-sectional, within-subject design. We report four main findings. First, unimanual tracking tasks assessed coordination of the dominant hand with a sensory, visual or passive-proprioceptive, signal. Young participants performed poorly and, despite continued improvement with age, performance was not equivalent to that of adults even in our oldest age groups, even if it was, at the same time, not credibly different. Second, symmetrical bimanual tasks assessed the ability to coordinate the two hands symmetrically both with and without visual hand position cues. Children performed these tasks with much less error than the unimanual tasks already at age 4. Accordingly, improvement across age was less marked, though, again, performance was not equivalent to that of adults even for the oldest age groups of our study. Moreover, bimanual coordination was independent of the visual display of hand position at all ages. Third, asymmetric motor tasks assessed the development of being able to overcome the motor symmetry bias through symmetric visual guidance. This ability was absent until age 9, and only at age 12 was asymmetric performance no longer credibly different from that of adults – and even at this age, performance was not completely adult-like, suggesting that further development would be observable throughout puberty. Fourth, evidence for cascade-like dependencies of the different functional aspects across development was weak. Whereas performance correlated highly within each set of tasks (unimanual, bimanual symmetric, 2:3 asymmetric), correlations were in the range of small to medium or entirely absent across these three functional domains. We will discuss these four points in turn.

### Unimanual tracking exhibits protracted and extensive development

Performance was comparable for visual and passive-proprioceptive tracking, and performance in the two tasks was highly correlated. These findings suggest a common developmental process for the use of sensory input for coordination. However, due to technical failure of our apparatus, the number of young participants in the proprioceptive task was low, which may have concealed differences especially in the affected, lower age groups.

On one side, the large variability and low accuracy of younger children in unimanual tracking strike as surprising, given that children learn to reach towards both external targets and their own body during their first year of life (Chinn et al., 2019; Somogyi et al., 2018) and adapt goal-directed hand movements to unexpected target displacement, visual rotations, and hand-diverting forces at age 4-5, (e.g. Contreras-Vidal et al., 2005; Ferrel-Chapus et al., 2002; Hay et al., 2005; Kagerer & Clark, 2014; Konczak et al., 2003). Such findings would seem to imply that sensorimotor control loops are established early on. Therefore it is critical to note that performance in sensorimotor adaptation improves until at least the age of 10 or 12 (e.g. Kagerer & Clark, 2014; P. H. Wilson & Hyde, 2013).

Similarly, somatosensory feedback is continually refined throughout development, evident, for instance, in increasingly reliable proprioceptive evaluation of arm position in arm matching tasks until late adolescence (Goble et al., 2005; Holst-Wolf et al., 2016). Our current finding that even the oldest children we tested did not quite perform adult-like is consistent with these prior reports. Children weight visual and haptic signals in a statistically optimal fashion in perceptive tasks at around 8-10 years of age (Gori et al., 2008). In our paradigms, children do not yet reach peak performance, suggesting additional developmental requirements in motor coordination, as compared to purely perception-based tasks. Indeed, other motor studies, too, have provided evidence for such extended developmental timelines (Goble et al., 2005; Martel et al., 2020). A critical difference with classical motor tasks is also that sensory information in adaptation tasks refers to the moving limb itself while the movement target remains fixed, whereas in the tasks used here, both the tracked signal and the tracking hand continuously change position. Furthermore, in experiments in which goal-directed movements are performed repeatedly, each new movement can be adapted according to the registered error of the previous movement(s). In contrast, continuous coordination requires continually integrating the external cue to derive the required correction of the ongoing movement. Our results suggest that these abilities develop later than those responsible for online corrections of discrete movements. Under this perspective, it is interesting to note that the developmental timeline of our tasks appears similar to that of target interception – another task that requires moving the body in response to a moving (rather than static) stimulus. Children around 12 can intercept moving objects similarly effectively as adults, even if some aspects of performance keep developing well into adolescence (Daum et al., 2007; Rothenberg-Cunningham & Newell, 2013).

### Symmetric bimanual coordination is motor-driven and available early, but still exhibits protracted development

Children in our study performed symmetric bimanual coordination with much higher accuracy than unimanual coordination already at age 4. Thus, availability of a motor command strongly improved coordination. This result fits with previous research that has demonstrated the general ability of children even at a young age to perform symmetrically, even if performance improves across childhood, likely in dependence of maturation of the corpus callosum (Cincotta & Ziemann, 2008; de Boer et al., 2012; Jeeves et al., 1988). The present results on uni-and symmetric bimanual coordination together suggest that the common motor command available in bimanual coordination is available early, well in line with the occurrence of uninstructed, mirror-symmetric movements in children (Cincotta & Ziemann, 2008; de Boer et al., 2012; Lazarus & Todor, 1987). In fact, availability of visual information on hand position did not improve circling performance at any age in our study, suggesting that symmetric bimanual coordination may be entirely motor-driven, at least for tasks like ours, in which positional information alone, but not spatial accuracy regarding the trajectory, is relevant. This finding supports previous work reporting that, in such tasks, visual feedback is not necessary or even detrimental (Fagard & Pezé, 1992). In contrast, integration was evident when children had to draw circles by hand within two boundaries, that is, a task that not only required positional symmetry, but also spatial accuracy in placing the pen (Lantero & Ringenbach, 2007) in contrast to simply pushing a handle forward, as in our apparatus. An influence of vision has also been demonstrated in other paradigms that involved self-guided, rather than spatially restricted, bimanual movements (Brandes et al., 2017; Mechsner et al., 2001). Thus, visual cues appear to be integrated only under specific, spatially challenging requirements.

Accordingly, symmetric bimanual coordination is quite evidently a different motor capability than unimanual coordination: given that the development of unimanual coordination lags several years behind that of bimanual coordination, it is unlikely that bimanual coordination is merely a special case of general coordinative abilities, as might be inferred from the suggestion that phase symmetry in motor coordination tasks such as the one we employed here is perceptually-driven (Bingham, 2004). It is all the more notable that, nonetheless, uni-and bimanual coordination were correlated in the range of 0.3 across all task pairs. Our correlation analysis controlled for age, so that the relationship between tasks does not simply reflect common improvement due to age. Thus, unimanual coordination with signals that are not self-produced appears to build partly on functionality that is implicit in symmetric bimanual coordination, although it cannot exploit a motor signal for coordination. What could be the common mechanism? Given that visual-motor integration did not appear relevant for our bimanual task, the link may be cognitive rather than sensory. Both types of tasks rely on continuous monitoring and relating two ongoing, constantly moving entities. In current models of motor control, monitoring is a feature of state estimation, that is, deriving the state of body and world based on issued motor commands, their expected sensory consequences, and the incoming sensory evidence. We suggest that it is refinement of such motor control functionality that is at the heart of the common development of our different tasks. In fact, development of state estimation functionality in the age interval investigated here has been previously suggested based on the finding that 6-8 year-old children estimated their hand position less precisely than older children and young adults (King et al., 2012).

### Asymmetric bimanual coordination is related to unimanual tracking

Children were first able to perform asymmetrical movement above chance level based on visual guidance in the 2:3 coordination task starting at age 9, and performance was better the older the child. These results are in line with previous studies that tested asymmetric coordination but employed Lissajous-type paradigms (Fagard, 1987; Fagard et al., 1985; Jeeves et al., 1988; Leinen et al., 2016). Asymmetric control of the hands requires uncoupling the two hemispheres’ motor commands, an ability which becomes increasingly efficient around age 10 when the corpus callosum matures, and other automatic, bilateral phenomena such as mirror movements subside (Fagard et al., 2001; Preilowski, 1972). Importantly, however, in our study performance in the 2:3 task was credibly correlated with the unimanual tasks. Thus, the ability to integrate visual feedback, as is required to track an external or passive-proprioceptive signal, is also important to use symmetric visual guidance to coordinate several limbs independently. In other words, visual guidance of motor control does not just rely on suppression of default hemispheric coupling, but also on the ability to couple with the sensory signal – and this ability, too, approached adult performance only around the age of 10 in our study. It is, moreover, of note that age 10 is also the time when children first exhibit adult-like optimal multisensory integration (Burr & Gori, 2012; Gori, 2015). Thus, multiple structural-anatomical, cognitive, and sensorimotor prerequisites appear to mature around the age of 10, together allowing the emergence of the perceptual guidance of movement (Bingham, 2004).

It is of interest that none of the participants succeeded in retaining the newly acquired coordination. This result implies that participants did not develop an internal model for the task’s motor structure, so that they were unable to correctly relate the consequences of their movement to the 2:3 phase requirements. Instead, they kept relying on visual feedback to correct their position online, which may explain the correlations with the unimanual tasks. We also find it worth pointing out that 2:3 coordination was unrelated to “regular”, symmetric bimanual coordination. One might have expected a negative correlation, with high performance in symmetric coordination presumably indicating particularly strong cross-hemispheric coupling or, alternatively, an inability to inhibit cross-hemisphere communication, respectively. However, such a relationship does not appear to exist.

### Evidence for non-linear development

We have so far emphasized the largely linear improvement across age in the different tasks devised in our study. In the 2:3 task, our analysis suggested that children may perform less successfully at age 8 than at ages 7 and 9. Notably, this interpretation rests on small numerical differences in our data and may, thus, not replicate. If taken at face value, this result suggests a non-linear effect in development, with a potential, temporary performance dip around age 8. Several previous studies have reported that children’s sensorimotor performance is temporarily impaired around the age of 8 (Badan et al., 2000; Hay, 1978; Martel et al., 2020; Pellizzer & Hauert, 1996; Smyth et al., 2004). Commonly, this temporary decline is interpreted to be due to a developmental change that involves employment of adult-like motor control strategies; this re-organization is thought to lead to transient disorganization of motor processing, with neither old nor new processes working with full efficiency.

Yet, even if our data support this notion of a transient performance decrement due to massive developmental restructuring, it is also salient that individual data points vary considerably. As is evident in our Fig. 3 and 5, which display single participant performance, many data points would not allow grouping a child into a specific age group with confidence. Rather, some very young children exhibit good performance, just as some much older children still exhibit rather poor performance. Whether this variability is simply indicative of interindividual differences in overall motor control, or rather of an extended time interval for development – with some children developing much faster than others – cannot be inferred from our cross-sectional data.

### Limitations

Clearly, our cross-sectional study design limits interpretation of our data not only in the afore-mentioned case. For instance, we cannot conclude whether all children undergo similar development but simply start at different time points, or whether development can be compressed or extended in some individuals. Similarly, the group-wide linear developmental trends do not imply similar, linear development in individuals. Instead, it is well possible that each individual exhibits step-like changes which cannot be uncovered when measuring every participant at only a single time point. Longitudinal assessment would, therefore, provide additional merit but requires exceptional financial and time-related effort. As is evident from our correlational analysis, the high number of participants of our study, together with the large number of experimental tasks, nevertheless allow valid conclusions about developmental questions.

A second shortcoming of our study is the undesirable loss we suffered due to technical failure. We lack data for almost half of the children in the unimanual, passive-proprioceptive tracking task, with especially high lack at the young age ranges.

Third, although we aimed at testing a wider-than-usual range of functions and tasks, we chose to focus on a single paradigm that involves the upper limbs. We deem the direct comparability of the 6 devised tasks a major strength of our study. Yet, this comparability comes at the price of unknowable generalizability to other paradigms and body parts.

Fourth, we have tried to estimate at which age the different tasks are probably mature by comparing children’s to adults’ performance. Such comparisons have caveats. For one, though differences between groups are not credible for some comparisons, the overall parameter probability distributions still look markedly different, and so one should not infer equivalence, especially given the sample size per age group. For practical equivalence, posterior distributions must be narrower to fit within the ROPE, which will require larger samples of same-age subgroups. Furthermore, it is commonly assumed that performance will remain adult-like once it has reached a respectively high performance level at some age; yet, transient impairment of some motor control functions has been reported during adolescence (Viel et al., 2009). Thus, characterization of performance in our six tasks at ages beyond those of our study awaits dedicated exploration.

Fifth, it would have been in the spirit of our overarching aim to obtain experimental data also from children above age 12. Our project funding and running time did not allow us to fulfill that wish. Development in the teen years strongly depends on the stage of puberty. It is therefore inadequate to simply compare teens by age, and we would have had to assess pubertal stage. Moreover, it is an open question how one would harmonize age-related data of young children with pubertal stage-related data from teens.

Finally, we have related some of our findings to what is known about callosal development, because the relationship of this structure with bimanual coordination is salient. Our study did not assess any neural measures and it is clear that many brain regions are involved in sensorimotor control (Boisgontier et al., 2018; Scott, 2012). Thus, we do not mean to imply that all of the findings we discuss depend solely on the corpus callosum.

### Summary and conclusions

To conclude, performance of children aged 4-12 across 6 experimental tasks based on a common, underlying movement coordination paradigm suggests interdependent, serial development of different motor control functions. Mastery of symmetrical bimanual coordination may rely predominately on motor-related, interhemispheric transfer and develops first. Yet, even this seemingly basic ability improves until at least age 9, likely dependent on callosal and cerebellar maturation. Unimanual synchronization with externally produced signals develops later and proceeds until at least age 11. Importantly, this ability depends in part on symmetric bimanual coordination, as suggested by correlations between bimanual and unimanual performance. This dependence may be mediated by their common reliance on state estimation for motor control. Finally, guiding asymmetric movement by focusing on visual symmetry develops last and appears to be based on multiple determinants, namely callosal maturation, multisensory integration, and flexibilization of the coupling between sensory and motor information.

## ACKNOWLEDGEMENTS

This work was supported by the German Research Foundation (DFG) through the Cluster of Excellence Cognitive Interaction Technology ‘CITEC’ at Bielefeld University, an Emmy Noether grant to TH (He 6368/1-1, 1-2, 1-3), and by the Collaborative Research Center SFB936 – 178316478 – B1/B11. We thank all children who participated in this study, and their parents. We thank Valerie Gockel, Alexander Seriyo and Frederick Thiemer for their support with data acquisition, Martin Bertele and Dennis Pauer for programming assistance, and Luke Miller, Yannish Ramgulam, Christian Seegelke, and Kenan Suljic for support with data analysis.

## DECLARATION OF CONFLICTING INTERESTS

There are no conflicts of interest.

## DATA AVAILABILITY STATEMENT

All data, code and material used in this study are available on the website of the Open Science Framework (OSF) at the following link: https://osf.io/x9u27/. Supplemental material can be found at https://osf.io/grnbx.

## Notes

https://osf.io/x9u27/

### Competing Interest Statement

The authors have declared no competing interest.

### Summary of Updates

New illustrations to improve readability, new analyses in the supplementary

