## Supplementaty Information for "Protracted development of visuo-proprioceptive integration for uni-and bimanual motor coordination"

### 1. Screening tests

Screening was based on parental information given via the Behavioral Rating Inventory of Executive Function (BRIEF) (Drechsler & Steinhausen, 2013) or the BRIEF-P for preschoolers (Daseking & Petermann, 2013). These questionnaires assign indices of behavior regulation, cognitive regulation, and overall executive function based on everyday life situations and are sensitive to attentional and learning disorders. We excluded eleven of the initial 138 children across all age groups because they scored above the standard threshold of one or more indices, indicative of atypical behavior.

Participants aged 7 and above were tested for IQ with the Zahlen-Verbindungs-Test (ZVT; English: number connection test or trail making test) (Oswald & Roth, 1978), in which children have to sequentially connect scrambled numbers on a piece of paper for 30/60 s (>10 yrs/<10 yrs of age). This test is recommended for fast screening in group testing, but not for individual correlation. We excluded three children who fell outside the 70-130 range. Because norm data do not exist below age 8, we applied norms for 8 year-olds also to the 7 year-olds; this was a conservative measure to ascertain a participant sample in the mean IQ range. IQ testing was, however, not possible for younger children, because they could not reliably read the numbers in the ZVT. Note, that we used IQ exclusively for participant selection, but did not include it as predictor in any of our statistical models.

### 2. Scenarios

Children could choose between 4 themes: Africa, Farm, Princess, and Sport. Each scenario was then adapted within the theme. Every two trials (1 or 2 sentences), the characters switched.

- Scenario example for Task 1 (Uni-Vis)

Hello! Today we are playing a catching game. Welcome to the farm! The dog must be as fast as the sheep. Are you ready? The cat is as fast as the dog. Now the sheep must be as fast as the cat. That was great. The animals have a lot of fun by running.

- Scenario example for Task 2 (Uni-Proprio)

Hello! Today we are playing a catching game. Welcome at the stadium. The boy shows you how fast the basketball must be. Now the two balls are rolling at the same speed. The body runs as fast as the ball to catch it. Well done!

- Scenario example for Task 3 (Bi-NoVis)

Hello! Today we are playing a catching game. Welcome in Africa. The elephant is gone. The monkey wants to catch him. Are you ready? The leopard also wants to run with the monkey. Now they are running together. Now the leopard and the elephant run together. Well done, the animals are having so much fun. Thank you for your help.

- Scenario example for Task 4 (Bi-Vis)

Hello! Today we are playing with balls. Welcome to the stadium. The boy practices with the basketball. Are you ready? You’re doing well, do not give up. The boy likes football very much and practices one more round. Well done! The boy is having so much fun. Thank you for your help.

- Scenario example for Task 5 (2:3-NoTransform)

Hello! Today we are playing a catching game. Welcome to the farm! The sheep and the dog are racing. But the dog is way faster. The dog and the cat are racing. But the cat is way faster. Now the cat and the sheep are running together. But the sheep is way faster. This is quite difficult, great how you held on!

- Scenario example for Task 6 (2:3- Transform)

Hello! Today the princess is dancing with the princes. Welcome to the castle. The frog-prince wants to be as fast as the princess. Are you ready? Remember that the prince is as fast. This is quite difficult, you can practice! Now the princess is dancing as fast as the prince. That was great. The princess and the frog-prince are dancing at the same speed. The dance continues. They have so much fun. Now the princess follows the prince one more time. It is difficult but you got this! The frog-prince is not tired yet, he follows the prince. Great, you are almost done. That was very long. The light is off, can you still help them? [This last sentence was the transition to 2-3:Retention (Task 7)].

### 3. Supplementary tables

**Table S1. Beta Estimates reflecting the comparison between adults and all other groups for Tasks 1 to 4 for the unsigned phase difference.** Positive β Estimates indicate that adults’ performance was better. Credible differences are indicated in bold (HDI outside [-3.9; 3.9]). ESS stands for the Effective Sample Size as calculated during Bayesian model fitting. The Lower and Upper Confidence Interval (CI) values constitute the 95% High Density Interval (HDI), which represent the interval that encompasses the 95% most probable values for a given parameter.

| **Tasks** | **Contrasts** | **β Estimate** | **Lower CI** | **Upper CI** | **ESS** |
| --- | --- | --- | --- | --- | --- |
| **Uni-Vis** | Adults vs. 12 yo | 9.6 | 0.6 | 18.9 | 13753 |
|  | Adults vs. 11 yo | 10.5 | 1.0 | 19.4 | 13170 |
|  | **Adults vs. 10 yo** | **18.2** | **10.1** | **26.8** | **12604** |
|  | **Adults vs. 9 yo** | **27.4** | **19.1** | **36.1** | **12516** |
|  | **Adults vs. 8 yo** | **42.8** | **33.7** | **52.1** | **13991** |
|  | **Adults vs. 7 yo** | **44.4** | **35.2** | **53.6** | **13544** |
|  | **Adults vs. 6 yo** | **55.3** | **46.2** | **64.4** | **13693** |
|  | **Adults vs. 5 yo** | **65.5** | **56.2** | **74.5** | **13231** |
|  | **Adults vs. 4 yo** | **73.1** | **64.2** | **82.0** | **13174** |
| **Uni-****Proprio** | Adults vs. 12 yo | 7.1 | -1.9 | 16.5 | 13726 |
|  | Adults vs. 11 yo | 9.3 | .17 | 18.5 | 14097 |
|  | **Adults vs. 10 yo** | **16.7** | **7.8** | **25.4** | **13815** |
|  | **Adults vs. 9 yo** | **26.3** | **17.2** | **34.8** | **13306** |
|  | **Adults vs. 8 yo** | **27.2** | **17.3** | **36.9** | **15359** |
|  | **Adults vs. 7 yo** | **26.6** | **16.9** | **36.1** | **14377** |
|  | **Adults vs. 6 yo** | **52.7** | **42.7** | **62.7** | **16132** |
|  | **Adults vs. 5 yo** | **72.8** | **60.6** | **84.7** | **19750** |
|  | **Adults vs. 4 yo** | **84.5** | **72.6** | **96.4** | **20715** |
| **Bi-NoVis** | Adults vs. 12 yo | 4.1 | -5.3 | 13.5 | 12723 |
|  | Adults vs. 11 yo | 3.9 | -5.4 | 13.4 | 12644 |
|  | Adults vs. 10 yo | 9.9 | 1.3 | 18.5 | 11733 |
|  | Adults vs. 9 yo | 11.7 | 3.1 | 20.4 | 11737 |
|  | **Adults vs. 8 yo** | **16.1** | **6.8** | **25.6** | **12938** |
|  | **Adults vs. 7 yo** | **16.6** | **7.1** | **26.1** | **12997** |
|  | **Adults vs. 6 yo** | **23.7** | **14.2** | **33.1** | **12951** |
|  | **Adults vs. 5 yo** | **33.8** | **24.3** | **43.3** | **12508** |
|  | **Adults vs. 4 yo** | **26.2** | **17.0** | **35.3** | **12582** |
| **Bi-Vis** | Adults vs. 12 yo | 5.3 | -4.0 | 14.7 | 13928 |
|  | Adults vs. 11 yo | 6.1 | -3.1 | 15.5 | 14250 |
|  | Adults vs. 10 yo | 12.1 | 3.8 | 20.9 | 13009 |
|  | **Adults vs. 9 yo** | **14.2** | **5.5** | **22.9** | **12949** |
|  | **Adults vs. 8 yo** | **19.2** | **9.8** | **28.5** | **14139** |
|  | **Adults vs. 7 yo** | **15.8** | **6.2** | **25.1** | **13722** |
|  | **Adults vs. 6 yo** | **23.1** | **13.8** | **32.3** | **14385** |
|  | **Adults vs. 5 yo** | **24.6** | **15.0** | **33.9** | **13747** |
|  | **Adults vs. 4 yo** | **29.6** | **20.6** | **38.7** | **13432** |

**Table S2. Beta Estimates reflecting the comparison between adults and all other groups for 2:3-NoTransform and 2:3-Transform for the unsigned phase difference.** Positive β Estimates indicate that adults’ performance was better. Credible differences are indicated in bold (HDI outside [-5.4; 5.4]). ESS stands for the Effective Sample Size as calculated during Bayesian model fitting. The Lower and Upper Confidence Interval (CI) values constitute the 95% High Density Interval (HDI), which represent the interval that encompasses the 95% most probable values for a given parameter.

| **Tasks** | **Contrasts** | **β Estimate** | **Lower CI** | **Upper CI** | **ESS** |
| --- | --- | --- | --- | --- | --- |
| **2:3-NoTransform** | Adults vs. 12 yo | -5.5 | -16.0 | 4.9 | 6475 |
|  | Adults vs. 11 yo | 8.0 | -2.2 | 18.4 | 6225 |
|  | Adults vs. 10 yo | 8.6 | -0.8 | 18.1 | 5838 |
|  | Adults vs. 9 yo | 10.8 | 1.2 | 20.3 | 6087 |
|  | Adults vs. 8 yo | 11.3 | .68 | 21.7 | 6733 |
|  | Adults vs. 7 yo | 12.5 | 2.1 | 22.9 | 6468 |
|  | Adults vs. 6 yo | 14.3 | 3.6 | 25.24.8 | 6537 |
|  | Adults vs. 5 yo | 15.5 | -0.5 | 31.9 | 12078 |
|  | Adults vs. 4 yo | 17.4 | 3.5 | 31.1 | 7915 |
| **2:3-Transform** | Adults vs. 12 yo | 9.3 | -1.0 | 19.5 | 7270 |
|  | **Adults vs. 11 yo** | **25.5** | **15.5** | **35.9** | **6754** |
|  | **Adults vs. 10 yo** | **34.9** | **25.4** | **44.31** | **6521** |
|  | **Adults vs. 9 yo** | **46.7** | **37.5** | **56.3** | **6838** |
|  | **Adults vs. 8 yo** | **47.5** | **37.0** | **57.7** | **7384** |
|  | **Adults vs. 7 yo** | **46.4** | **36.2** | **56.7** | **7068** |
|  | **Adults vs. 6 yo** | **60.3** | **49.8** | **70.6** | **7374** |
|  | **Adults vs. 5 yo** | **59.4** | **43.2** | **75.8** | **12986** |
|  | **Adults vs. 4 yo** | **63.7** | **50.3** | **77.1** | **8685** |

### 4. Control task: Spatial span

This classic test to assess visuospatial working memory (Berch, Krikorian, & Huha, 1998) has been adapted by BKIN technologies on the Kinarm. For each trial, participants had to reproduce a sequence of square that have been randomly lighten up, by reaching and pausing on the specific squares. Twelve squares were displayed in a 3 x 4 grid. The first sequence comprised 4 squares, and the next sequence then had 1 square more when the previous sequence was correct, or 1 square less in case of an error in the previous sequence. There were 2 practice trials followed by 16 analyzed trials. The younger children (4-5 yo) could rarely reach for the last row in the standardized task setup, resulting in abortion of the task after a few trials. Duration: about 3min.

Data on spatial span was available on 112 of the 132 subjects, including the 12 adults. We extracted the mean spatial span of each participant as the mean number of squares they could remember correctly for each trial. Data is plotted in Fig. S1.

Participants performed as expected in the spatial span task, inspired from the Corsi block. It has indeed been reported that around 8 years, children scored 4 in the original task, as they did here (Roden, Kreutz, & Bongard, 2012). Additionally, a recent study tested the span capacity of adolescents from 11 to 20 years in a similar Corsi-inspired task (Burggraaf, Frens, Hooge, & Geest, 2018). They scored between 3 and 7, which matches our present observations. Previously it has been shown that children reached their adult performance during early adolescence, usually around 14 years (Farrell Pagulayan, Busch, Medina, Bartok, & Krikorian, 2006). Here it seems that the 12 year-olds already exhibited similar performance to adults.

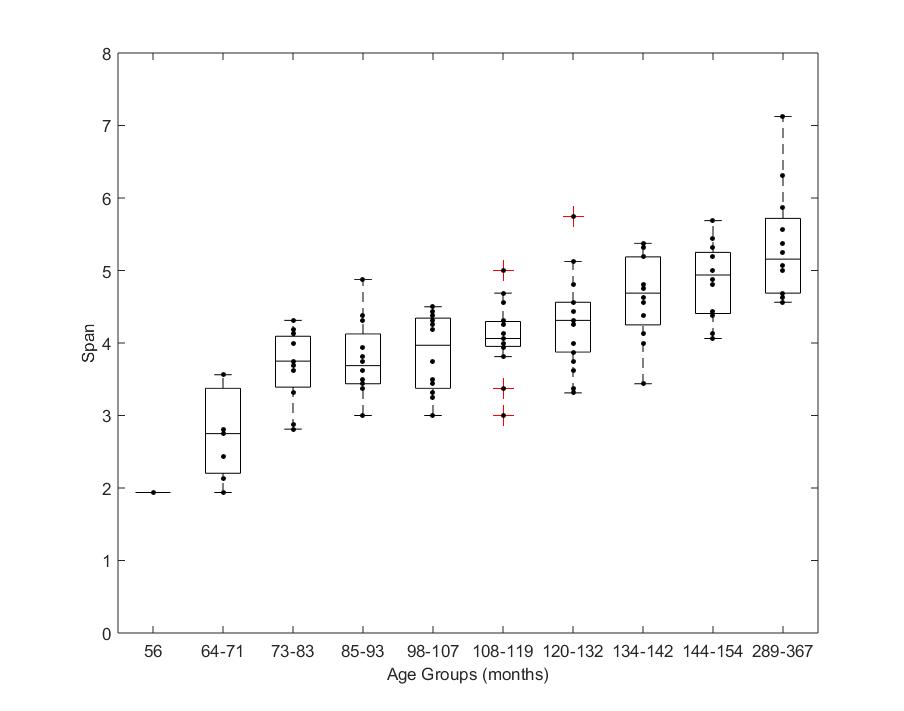

**Figure S1. Spatial span as a function of age, for each age group.** Numbers indicate the age range. Only one 4-year-old child (age = 56 months) was able to complete the task.

### 5. Percentage of adequate coordination ≈ proportion on time-on-task

The mean proportion of the task during which participants maintained the correct coordination (± 30°) increased with age. Below a table of these means for each task (Table S4). Note that by doing a 1:1 coordination in the 2:3 tasks, the percent of random correct coordination should be around 24%, very close to what we obtain. The 2:3-Transform task was constituted of three blocks. These three blocks had very similar scores, indicating that whatever was the initial performance, it did not improve within the 10 minutes of the task.

**Table S3. Percent of the task within the correct coordination for each group.**

|  | **4 yo** | **5 yo** | **6 yo** | **7 yo** | **8 yo** | **9 yo** | **10 yo** | **11 yo** | **12 yo** | **Adults** |
| --- | --- | --- | --- | --- | --- | --- | --- | --- | --- | --- |
| **Uni-Vis** | 17 | 19 | 24 | 28 | 26 | 40 | 47 | 58 | 56 | 72 |
| **Uni-Proprio** | 9 | 14 | 20 | 43 | 55 | 37 | 42 | 58 | 62 | 73 |
| **Bi-NoVis** | 59 | 53 | 65 | 71 | 70 | 73 | 75 | 83 | 86 | 93 |
| **Bi-Vis** | 62 | 59 | 68 | 71 | 69 | 72 | 78 | 84 | 87 | 94 |
| **2:3-NoTransform** | 22 | 22 | 25 | 23 | 24 | 23 | 24 | 27 | 35 | 30 |
| **2:3-Transform - 1° Block** | 22 | 25 | 21 | 27 | 29 | 28 | 37 | 41 | 54 | 65 |
| **2:3-Transform - 2° Block** | 18 | 21 | 23 | 30 | 30 | 30 | 37 | 43 | 51 | 64 |
| **2:3-Transform - 3° Block** | 21 | 21 | 24 | 27 | 30 | 30 | 37 | 45 | 55 | 61 |
| **2:3-Retention** | 15 | 20 | 16 | 18 | 16 | 18 | 18 | 23 | 30 | 29 |

### 6. Signed phase difference of Uni- and bimanual, symmetric coordination (Tasks 1-4)

We assessed whether one hand had a constant offset relative to the other, that is, whether one hand regularly leads the other by a few degrees in a model that predicted the signed phase difference between the two coordinated signals – a visual or proprioceptive signal and a hand (unimanual tasks), or the two hands (bimanual tasks). Figure S2 illustrates performance in the four tasks included in the model.

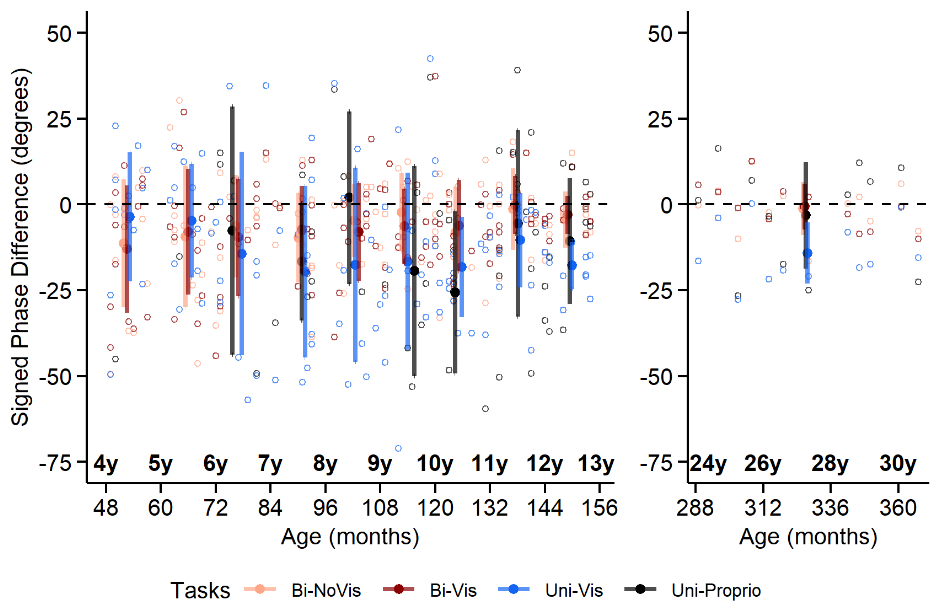

**Figure S2. Signed phase difference between the two hands in unimanual Tasks 1 and 2 (blue and black) and bimanual Tasks 3 and 4 (dark and light red) for children (left panel) and adults (right panel).** The dashed lines represent the 0° threshold: values above this threshold indicate that the dominant hand is leading, values below indicate that the dominant hand is lagging. Colored circles represent the mean for each age group; bars show the s.d. Fewer than half of the 4-8 year-old participants performed Task 2 (Uni- Proprio; black), the means are thus likely to be non-representative (means in Uni-Proprio are not shown for 4- to 5-year-old children, because n ≤ 2).

In the two unimanual tasks, participants had to synchronize the movement of their dominant hand with a circling dot (Uni-Vis) or with their passively moved, non-dominant hand (Uni-Proprio). If a participant moves the hand following an external guiding signal, there should be a constant offset between the guiding signal and the hand, given that processing of sensory information takes time. If, in contrast, the hand maintained a zero phase lag, this would imply that the hand is moved predictively. As can be seen from Figure S2 and Table S4, the intercept for both unimanual tasks was credibly different from zero, but with no improvement with age. There was no difference between the tasks, suggesting that the dominant hand lagged behind the non-dominant hand for all ages and that participants did not anticipate the movement of the dot/passively moved arm.

Regarding the bimanual tasks, where participants had to coordinate both hands, the intercept was not credibly different from zero (HDI_95%_ overlapped with the ROPE [-5.1; 5.1]), and there was no improvement with age in line with the idea that there was no leading/lagging hand.

**Table S4.** **Beta Estimates reflecting the comparison between Tasks 1 to 4 in Model 1**. For Intercept, negative β Estimates indicate that the dominant hand lagged behind the non-dominant hand. For Age, negative β Estimates indicate that performance was better with higher age (smaller error for each standardized unit of Age of 30.3 months, see Methods). For Task, positive β Estimates indicate that Intercept was higher in the first task of each line, meaning that for this specific task, there was a higher error at younger ages. The interaction Task × Age reflects differences in steepness of improvements with Age. The Lower and Upper Confidence Interval (CI) values constitute the 95% High Density Interval (HDI), which represent the interval in which 95% of the values will be found. Credible differences are indicated in bold (HDI outside of the Region Of Practical Equivalence in which there is no effect [-5.1; 5.1]). ESS stands for the Effective Sample Size as calculated during Bayesian model fitting.

| **Effect** | **Contrasts** | **β Estimate** | **Lower CI** | **Upper CI** | **ESS** |
| --- | --- | --- | --- | --- | --- |
| **Intercept** | **Uni-Vis** | **-14.2** | **-16.9** | **-11.8** | **6102** |
|  | **Uni-Proprio** | **-13.5** | **-16.5** | **-10.5** | **7995** |
|  | Bi-NoVis | -6.4 | -9.0 | -3.8 | 6133 |
|  | Bi-Vis | -6.3 | -8.9 | -3.8 | 6257 |
| **Age** | Uni-Vis | -3.5 | -5.9 | -1.9 | 6786 |
|  | Uni-Proprio | 1.7 | -1.3 | 4.6 | 9706 |
|  | Bi-NoVis | 2.62 | 0.1 | 5.2 | 6850 |
|  | Bi-Vis | 3.1 | .55 | 5.6 | 7416 |
| **Task** | Uni-Proprio – Uni-Vis | .76 | -1.5 | 3.0 | 56949 |
|  | Bi-Vis – Bi-NoVis | 0.1 | -1.7 | 1.8 | 49138 |
|  | **Uni-Vis – Bi-NoVis** | **-7.9** | **-9.5** | **-6.3** | **47591** |
|  | **Uni-Vis – Bi-Vis** | **-8.0** | **-9.5** | **-6.4** | **67683** |
|  | Uni-Proprio – Bi-NoVis | -7.1 | -9.5 | -4.8 | 31485 |
|  | Uni-Proprio – Bi-Vis | -7.2 | -9.5 | -4.9 | 59351 |
| **Interaction**  **Task × Age** | Uni-Proprio – Uni-Vis | 5.2 | 2.9 | 7.5 | 56290 |
|  | Bi-Vis – Bi-NoVis | .61 | -1.2 | 2.4 | 33806 |
|  | Uni-Vis – Bi-NoVis | -6.1 | -7.6 | -4.5 | 33588 |
|  | Uni-Vis – Bi-Vis | -6.6 | -8.2 | -5.0 | 64376 |
|  | Uni-Proprio – Bi-NoVis | -0.9 | -3.3 | 1.5 | 34199 |
|  | Uni-Proprio – Bi-Vis | -1.4 | -3.7 | 1.0 | 55085 |

Given that there was no indication for a systematic trend dependent on age, we do not report by-year-of-age group analyses.

Moreover, we did not assess the signed difference for the 2:3 coordination, because the instructions were to be faster with one hand, and whether one or the other transformed cursor leads the other would not be informative about general aspects of coordination.

### 7. Sensory dominance in bimanual motor coordination: Retention task

In Task 7, 2:3-Retention, participants had to concurrently perform three circles with their dominant, and two with their non-dominant hand. Each trial began just like 2:3-Transform, that is, with transformed visual feedback. However, after 5 s, the visual feedback was removed. Participants had to continue the 2:3 coordination, allowing us to assess their ability to perform the coordination without online feedback after having previously learned the task before.

Sample size in each group for this task:

| **Age Group** | **4 yo** | **5 yo** | **6 yo** | **7 yo** | **8 yo** | **9 yo** | **10 yo** | **11 yo** | **12 yo** | **Adults** |
| --- | --- | --- | --- | --- | --- | --- | --- | --- | --- | --- |
| **Number of included participants** | **n = 14** | **n = 12** | **n = 12** | **n = 12** | **n = 12** | **n = 16** | **n = 18** | **n = 12** | **n = 12** | **n = 12** |
| **Task 7 – 2:3-Retention** | 3 | 1 | 7 | 11 | 11 | 16 | 17 | 12 | 12 | 12 |

To investigate the ability of children to learn an asymmetric coordination, the 2:3 tasks were analyzed with a Bayesian mixed model. Below is the report of 2:3-Retention. The standardized unit of age was 24.6 months, with a ROPE of [-5.4; 5.4].

Performance did not improve with age (β_Age_ = -4.9, HDI_95%_ = [-7.1; -2.6]). There was a credible difference for both *Task* and *Task* × *Age* interaction contrasts, indicating that the intercept and rate of improvement was higher for 2:3 Transform than 2:3-Retention (β_Task_ = 18.1, HDI_95%_ = [16.8; 19.4]; β_Interaction_ = 8.5, HDI_95%_ = [7.1; 9.9]). By contrast, there was no credible difference between the 2:3-Retention and 2:3-NoTransform (β_Task_ = 4.0, HDI_95%_ = [2.7; 5.4]; β_Interaction_ = -0.8, HDI_95%_ = [-2.3; 0.6]) indicating that the transformation was not learned without constant feedback.

As there was no indication of any performance change in dependence of age, we do not report by-year-of-age group analyses.
